# Multicolored, Sonosensitizer-optimized Organic Mechanoluminescent Nanoparticles for Functional Sono-optogenetics

**DOI:** 10.64898/2025.12.16.694777

**Authors:** Xiangping Liu, Wenliang Wang, Brinkley Artman, Jiefeng Diao, Yangyang Zhao, Weilong He, Sihan Yu, Kai Wing Kevin Tang, Mengmeng Yao, Christina Gu, Brian Song, Huiliang Wang

**Affiliations:** Biomedical Engineering Cockrell School of Engineering, The University of Texas at Austin, Austin, Texas 78712, United States; Department of Chemistry, The University of Texas at Austin, Austin, TX 78712, United States; Department of Neuroscience, The University of Texas at Austin, Austin, Texas 78712, United States

**Keywords:** Ultrasound, sonosensitizers, ROS, mechanoluminescence, sono-optogenetics

## Abstract

Light enables precise visualization and control of cellular processes, but its utility in deep tissues is fundamentally limited by poor optical penetration, particularly in the deep brain. Ultrasound-triggered mechanoluminescence offers a non-invasive strategy for remote light delivery, yet existing organic systems remain monochromatic and low-intensity, largely due to an incomplete understanding of ultrasound-induced emission. Here, we report a multicolor mechanoluminescence platform that couples reactive oxygen species-responsive chemiluminescent donors with fluorescent acceptors via Förster resonance energy transfer, generating tunable emission from blue (459 nm) to red (592 nm). Systematic screening potentially reveals that electronic energy gap–dependent reactive oxygen species generation serves as a predictive design principle for high-performance mechanoluminescent materials. The emitted spectrum and intensity are sufficient to activate ChR2 and ChRmine and inhibit eOPN3, enabling bidirectional, fiber-free neuromodulation under focused ultrasound. By integrating spatially precise ultrasound with programmable photon output, this platform establishes a non-invasive strategy for deep-tissue neural monitoring and provides a foundation for applications in bioimaging, gene editing, and precision therapeutics.

## Introduction

Light plays a central role in monitoring cellular events and deconstructing complex organismal behaviors.^1^ Organisms have independently evolved specialized organs and proteins to sense light across various wavelengths and intensities, enabling functions such as vision, circadian rhythm regulation, and other critical physiological and sensory processes.^2,3^ In recent decades, light-based technologies, including optogenetics,^4^ molecular imaging,^5,6^ and light-activated gene editing,^7^ have revolutionized our ability to visualize and manipulate dynamic cellular processes in vivo, substantially advancing our understanding of pathophysiology and driving the development of innovative therapeutic strategies.^8^ However, their efficacies are limited by insufficient light penetration due to increasing scattering and absorption that scale with depth. While recent advancements in optical transparency technologies have enhanced depth-light delivery capabilities, invasive fiber implantations are usually required to achieve remote light delivery in deep tissues.^9^ These constraints highlight the pressing need for non-invasive strategies to remotely deliver light into tissues, a critical step toward advancing light-based technologies to the next stage of clinical and research utility.

Ultrasound-triggered mechanoluminescence (ML) presents an exciting opportunity for remote and non-invasive light delivery in deep tissue with high spatiotemporal resolution and excellent safety.^10^ Piezoelectric nanocrystals have been developed as internal light sources capable of converting mechanical energy from ultrasound into light, facilitating deep-tissue optogenetic stimulation termed sono-optogenetics.^11^ However, the applicability of these inorganic nanomaterials is limited by their need for recharging, poor light emission repeatability, and potential toxicity due to heavy metal content. Organic mechanoluminescent materials, particularly those based on luminol and 1,2-dioxetane derivatives, offer a biocompatible alternative. These systems exploit reactive oxygen species (ROS) generated through ultrasound-induced cavitation to initiate chemiluminescence, enabling non-invasive imaging and therapy.^12^ While great progress has been made in ultrasound-triggered organic mechanoluminescent materials, existing systems typically exhibit monochromatic emission and low luminescent intensity, largely due to incomplete understanding of the underlying mechanisms governing ultrasound-induced ROS production and energy transfer.^13,14^ Thus, the development of high-efficiency, multicolor, and tunable sono-mechanoluminescence remains a significant challenge.

In this study, we report on a multicolor mechanoluminescent platform that integrates ROS-triggering chemiluminescent donors with a library of fluorescent acceptors via Förster resonance energy transfer (FRET). By optimizing donor-to-acceptor ratios, we achieve tunable light emissions spanning from blue (459 nm) to red (592 nm). Moreover, we systematically screen a panel of candidate sonosensitizers and identify a strong correlation between their electronic energy gaps and ROS generation efficiency, establishing a predictive strategy for designing next-generation mechanoluminescent materials with enhanced performance. To demonstrate functional applicability, we show that the emission spectrum of ultrasound-triggered ML encompasses the activation windows of quintessential optogenetic actuators, including ChR2 (blue), eOPN3 (green), and ChRmine (red), and that the emitted intensity is sufficient to drive both excitatory and inhibitory neuromodulation under focused ultrasound. We anticipate that this work will inspire the development of mechanoluminescent materials as a versatile platform for non-invasive light-based technologies beyond sono-optogenetics, including applications in bioimaging, gene editing, and other precision therapeutics.

**Scheme 1.**
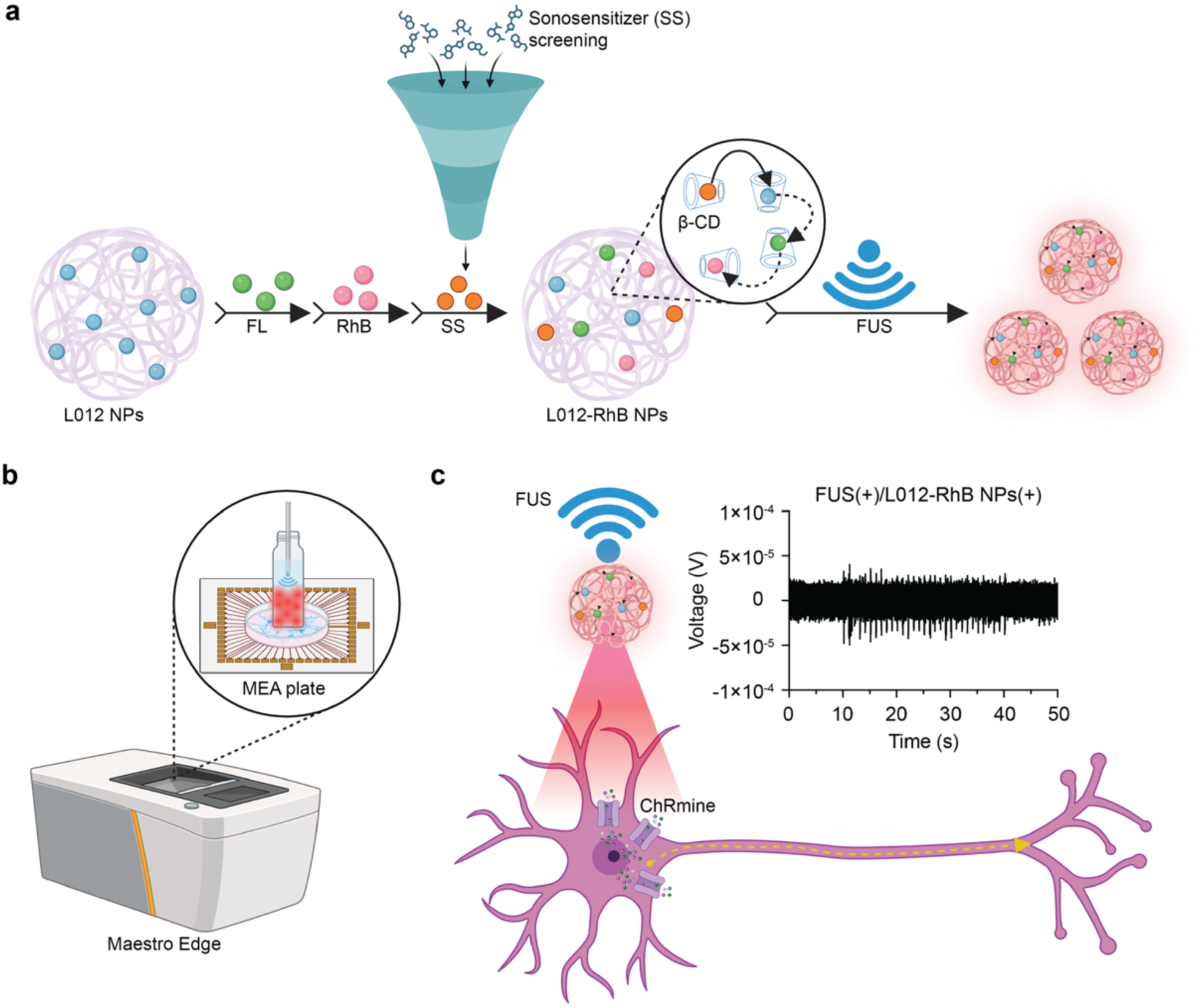
Schematic illustration of the design of a multicolor mechanoluminescent nanoparticle system (exemplified by L012-RhB NPs) and its application in sono-optogenetic activation of neurons expressing ChRmine, created via BioRender.com. (**a**) Mechanisms of nanoparticle fabrication and luminescence emission by L012-RhB NPs under ultrasound stimulation. (**b**) Illustration of ultrasound-triggered luminescence from L012-RhB NPs and the resulting sono-optogenetic stimulation of cultured primary neurons as recorded by a microelectrode array. (**c**) After neuronal transduction with ChRmine via adeno-associated virus (AAV), L012-RhB NPs are activated by FUS to emit red light, subsequently triggering ChRmine, with neuronal electrical signals recorded through macroelectrode arrays shown in the inset.

## Results and Discussion

### FRET-Based Multicolored Chemiluminescent Systems

To enable tunable light emission, we first synthesized blue light-emitting nanoparticles (L012-NPs) as the primary energy donor and evaluated their chemiluminescent properties. L012-NPs were prepared by loading L012, Tris base, and cobalt (II) ions into β-CD to form dynamic nanoassemblies above the critical concentration of β-CD, fulfilling roles as chemiluminescent molecules, stabilizing and intermediate-binding agents, and Lewis-acid catalysts, respectively.^15,16^ L012 acts as an ROS-reactive chemiluminescent substrate. In the presence of H_2_O_2_, L012 is activated, yielding L012-NPs with robust blue chemiluminescence at an emission peak of 459 nm (**Figure S1**). Next, we prepare chemiluminescent nanoparticles with different wavelengths by modulating the components of the FRET. Green emission (524 nm) was first achieved by incorporating the fluorophore fluorescein (FL) into L012-NPs (L012-FL NPs, **Figure 1a**), where the energy transfers from excited L012 donor to an FL energy receptor, thus emitting green light. FRET efficiency is dominated by the ratio between donors and receptors. Therefore, to optimize FRET efficiency and maximize luminescence intensity, FL concentrations ranging from 1.0 mM to 2.0 mM (in increments of 0.2 mM) are evaluated (**Figure 1b-d**). The results show that increasing FL concentration resulted in gradual decreases in individual spectra of L012 and FL, with an optimal FRET efficiency of 95% achieved at 1.4 mM (**Figure 1d**), and maximum accumulated chemiluminescence intensity is observed at 1.2 mM FL concentration (**Figure 1b and Figure 1c**). After that, we also expect to achieve longer-wavelength emissions, and secondary fluorophores are introduced into the L012-FL nanoparticle systems for on-demand light emission through cascade FRET (**Figure 1e-k**). Our results demonstrate that 550 nm, 573 nm and 592 nm chemiluminescence is specifically generated as the introduction to a secondary receptor fluorophore, including eosin Y (EY), phloxine B (PB), and RhB, where the FRET efficiency was optimized up to 85% at 0.2 mM EY (**Figure 1i**), 75% at 1.2 mM PB (**Figure 1j**) and 62% at 1.2 mM RhB (**Figure 1k**). Optical imaging and accumulated chemiluminescence intensity measurements of nanoparticles containing EY (**Figure 1e**), PB (**Figure 1f**), and RhB (**Figure 1g**) also confirmed expected color profiles, although overall intensity decreased with higher fluorophore concentrations (**Figure 1h**). By employing single-step and cascade FRET strategies, we successfully developed a suite of nanoparticles emitting across a spectrum from blue to red, spanning approximately 130 nm (**Figure 1l**). Dynamic light scattering (DLS) analysis confirmed nanoparticle hydrodynamic diameters averaging approximately 180 nm, with negative zeta potentials (**Figure 1m**).^17^ Detailed experimental data, including nanoparticle composition, molar ratios, size, PDI, zeta potential and emission peaks, are summarized in **Figure 1n**.

**Figure 1.**
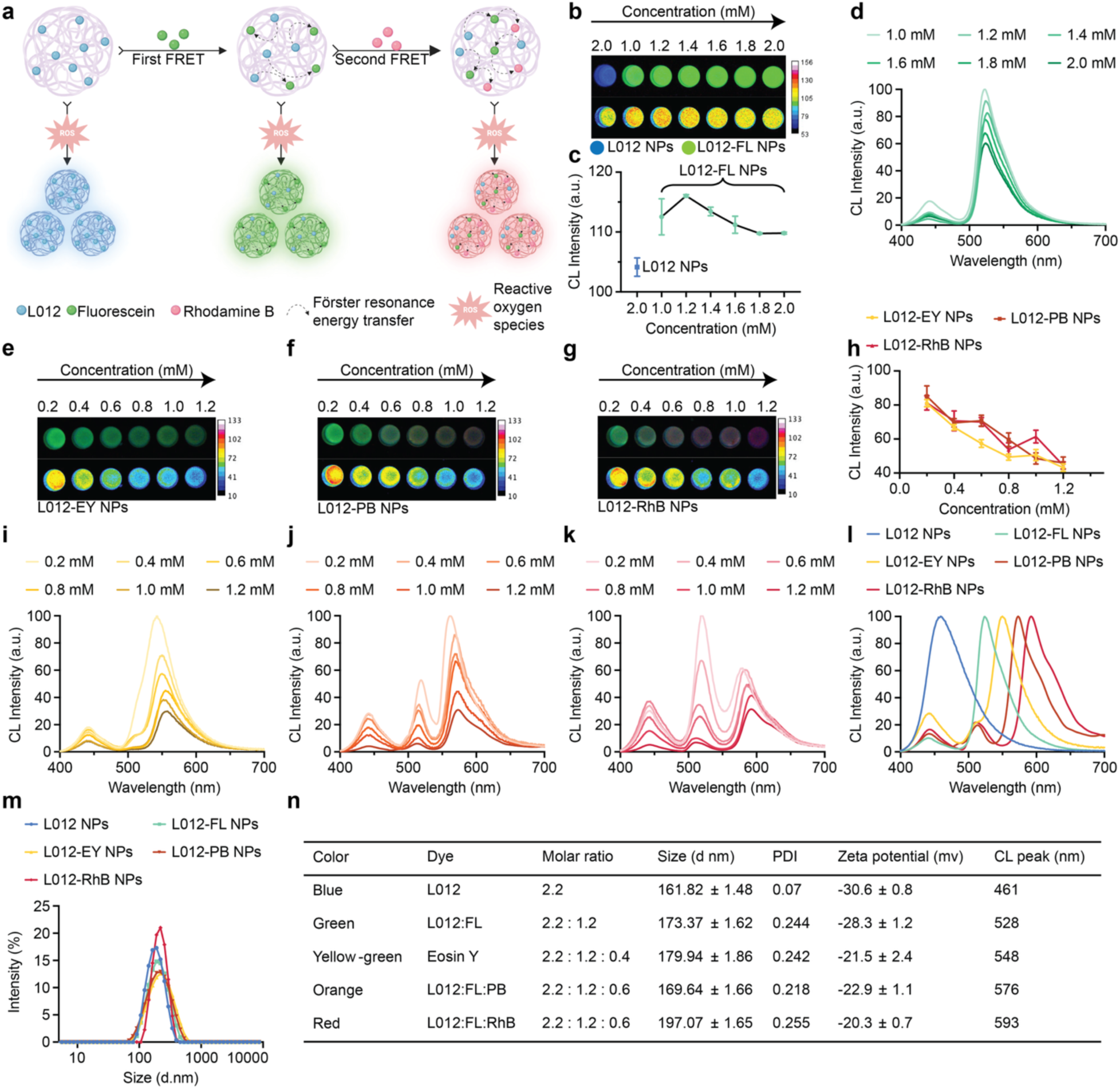
Chemiluminescence (CL) characterization and optimization of multicolor L012-based nanoparticles. (**a**) Construction schematic of multicolor nanoparticles employing highly efficient single and cascade FRET, wherein L012 acts as donor and FL, EY, PB, or RhB serve as acceptors for achieving progressively longer wavelengths, created via BioRender.com. (**b-c**) Optical images and intensity quantification of L012 NPs and L012-FL NPs under H_2_O_2_. (**d**) CL spectra of varying FL concentrations in L012-FL NPs. (**e-g**) Optical images of L012-EY (**e**), L012-PB (**f**), and L012-RhB (**g**) nanoparticles at different concentrations. (**h**) Concentration-dependent CL intensities of L012-EY, L012-PB and L012-RhB NPs. (**i-k**) CL spectra of L012-FL NPs incorporating increasing concentrations of EY (**i**), PB (**j**), and RhB (**k**), respectively. (**l**) CL spectra displaying five distinct emission colors. (**m**) Size distribution profiles of these nanoparticles via DLS. (**n**) Compositional and physicochemical characterization of the multicolor nanoparticle system.

### Multicolored Mechanoluminescent Systems

After optimizing the chemiluminescent properties of the L012-based nanoparticles, we further investigated their potential for ultrasound-controllable light emissions for optical delivery. As an efficient ROS scavenger, L012 could sense the ROS and be activated for light emission. Thus, IR780, a potent sonosensitizer proven to efficiently generate ROS, particularly ^1^O_2_ and •OH (**Figure 2a**) is integrated into L012-based nanoparticles.^18–21^ Upon ultrasound stimulation, these nanoparticles exhibited robust and pulse-dependent light emissions, clearly visible in the optical images captured during ultrasound-off and ultrasound-on states (**Figures 2b–2g**). Notably, Sono-mechanoluminescent (Sono-ML) intensity remarkably decreased as emission wavelengths became longer (**Figures 2h**). This reduction probably results from the substantially shorter lifetimes of ultrasound-generated ROS (approximately 3 µs for ^1^O_2_ and 1 ns for •OH),^22,23^ compared to the prolonged stability of H_2_O_2_ used traditionally in chemiluminescence, allowing efficient FRET processes. We further evaluated Sono-ML intensity under varying ultrasound frequencies (**Figure 2i**) and durations (**Figure 2j**). Consistent and highly synchronized Sono-ML responses were observed at frequencies of 1, 2, 4, and 8 Hz, demonstrating rapid and stable photon emission, essential for effective neuronal activation in sono-optogenetics. Moreover, consistent Sono-ML intensities under ultrasound pulses ranging from 100 ms to 900 ms underscore the suitability of these nanoparticles for sustained neuromodulation. ML intensity was also evaluated across a range of ultrasound peak pressures. Minimal luminescence was observed at 0.71 MPa, with ML intensity progressively increasing at higher pressures (0.71-1.67 MPa), a pattern consistently observed across all nanoparticle variants (**Figure 2k**). The total effective ML duration, assessed under a cyclic ultrasound regime of 1 second on/off, demonstrated luminescence sustainability for approximately 15 minutes, adequate for real-time bioimaging applications (**Figure 2l**). Comparative analysis revealed that our β-CD@IR780/L012 nanoparticles (L012 NPs) exhibit significantly longer luminescence durations, 5.4 and 2.5 times longer than previously reported Lipo@IR780/L012 and HOF@L012 systems, respectively (**Figure 2m**).^13,24^ These comprehensive findings demonstrate that our organic mechanoluminescent systems provide bright, repeatable, tunable, stable, and durable luminescence, holding significant promise for various biomedical applications, including sono-optogenetics, bioimaging, and theranostics.

**Figure 2.**
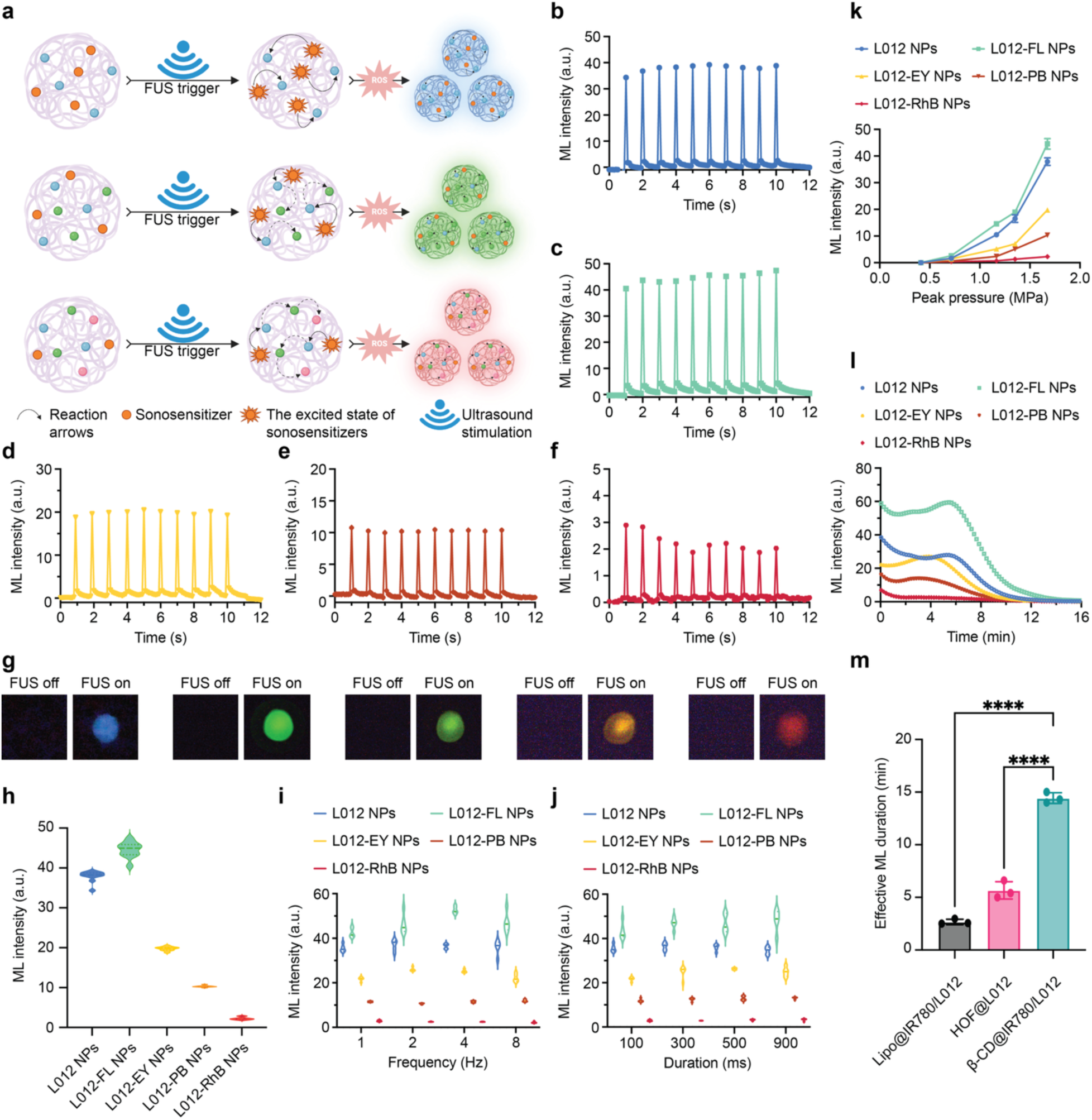
Sono-ML characterization of multicolor L012-based nanoparticles. (**a**) Schematic representation of the multicolor Sono-ML emission mechanism for L012-based nanoparticles under FUS stimulation, created via BioRender.com. (**b-f**) The dynamic intensity profiles of L012, L012-FL, L012-EY, L012-PB and L012-RhB NPs emissions under intermittent ultrasound irradiation (1.5 MHz, 1.55 MPa, 100 ms on, 900 ms off). (**g**) True-color emission optical images of these nanoparticles under the same ultrasound irradiation. (**h**) Comparative ML intensities of all nanoparticles under standardized conditions. (**i-j**) ML intensities of the multicolor nanoparticle platforms at various ultrasound frequencies and pulse intervals (1.5 MHz, 1.55 MPa). (**k**) ML intensities of all nanoparticles at increasing ultrasound peak pressures. (**l**) Effective ML durations for all nanoparticle variants under cyclic FUS stimulation. (**m**) Comparative ML duration analysis of Lipo@IR780/L012, HOF@L012, and β-CD@IR780/L012.

### Understanding Ultrasound Activation of Sono-sensitizers for Sono-mechanoluminescence

The intensity of Sono-ML generated by L012-based nanoparticles strongly correlates with the efficiency of ROS generation by sonosensitizers under ultrasound stimulation.^14,25^ To optimize Sono-ML intensity and guide the design of Sono-ML systems with better performance, we systematically evaluate common small organic sonosensitizers categorized into porphyrin, cyanine, and xanthene groups based on their chemical core structures (**Figure 3a**).^26–29^ Mechanoluminescence from the Sono-ML system depends strongly on sonosensitizer concentration, with an optimum at around 0.1 mg/mL (**Figure S2**). We assess ^1^O_2_ generation efficiency using the well-established scavenger 1,3-diphenylisobenzofuran (DPBF).^30^ DPBF reacts specifically with ^1^O_2_ via cycloaddition and decomposition, yielding 1,2-dibenzoylbenzene (**Figure 3b**), which possesses a prominent absorption peak at 420 nm. •OH production is evaluated via the degradation of methylene blue (MB), which has a characteristic absorption peak at 680 nm and transforms into benzenesulfonic acid and 4-nitrophenol upon reaction with •OH (**Figure 3c**).^31^ Taking indocyanine green (ICG) as an example, we exposed the ICG-DPBF solution to ultrasound transducer and found minimal spectral changes without ultrasound irradiation (**Figure 3d**). Upon ultrasound stimulation, the DPBF absorption peak at 420 nm decreased over time, confirming the generation of ^1^O_2_. Notably, ICG degraded faster than DPBF, primarily due to its relatively small HOMO-LUMO energy gap, facilitating energy transfer from ultrasound-induced mechanical stress (**Figure 3e**).^32^ Quantitative analysis revealed approximately 45% DPBF degradation after 90 s of FUS, compared to less than 7% without ultrasound (**Figure 3f**). Similar control experiments with ICG-MB showed negligible spectral changes without ultrasound stimulation (**Figure 3g**), while significant MB absorption decrease was observed with ultrasound (**Figure 3h**). Quantitative analysis demonstrated 38% MB degradation after 90 s of FUS irradiation, compared to 12% without ultrasound due to natural MB aggregation and decomposition in an aqueous environment (**Figure 3i**).^33^ We further measured ROS generation efficiency for fourteen additional sonosensitizers (**Figure S3 to Figure S16**). Cyanine derivatives exhibited relatively lower ^1^O_2_ generation, whereas xanthene derivatives demonstrated notably higher efficiency, likely due to the heavy-atom effect enhancing intersystem crossing (ISC) via spin-orbit coupling (SOC), increasing triplet excited-state yield (**Figure 3j**).^34–36^ However, •OH production efficiencies were generally lower and similar across all sonosensitizers, likely because •OH generation primarily relies on ultrasound-induced pyrolysis rather than molecular structure alone (**Figure 3k**).^37,38^ To elucidate the underlying mechanism of ROS generation, we explored correlations between ROS yields and various electronic energy gaps using Jablonski diagrams.^39,40^ We hypothesized that the generation efficiency of ^1^O_2_ primarily depends on the S_1_-S_0_ energy gap, whereas •OH production efficiency relates to the HOMO-LUMO gap (**Figure 3l**). A larger S_1_-S_0_ gap exponentially suppresses the competing internal conversion pathway. This increases the S_1_ lifetime, maximizing the quantum yield of ISC and thus T_1_ population.^41–43^ A smaller HOMO-LUMO gap indicates higher intrinsic redox reactivity. This enhanced reactivity is the definition of the Type I mechanism, facilitating the electron transfer cascade to produce radicals including •OH.^44–46^ Indeed, the positive correlation (Pearson correlation r = 0.73) observed between relative ^1^O_2_ generation efficiency and S_1_-S_0_ energy gap enabled the establishment of a predictive model for ^1^O_2_ yields from untested sonosensitizers based on calculated energy gaps (**Figure 3m**). Likewise, a negative correlation with Pearson correlation r of −0.41 between relative •OH generation efficiency and the HOMO-LUMO energy gap was also established, allowing prediction of •OH yields from additional sonosensitizers (**Figure 3n**).

**Figure 3.**
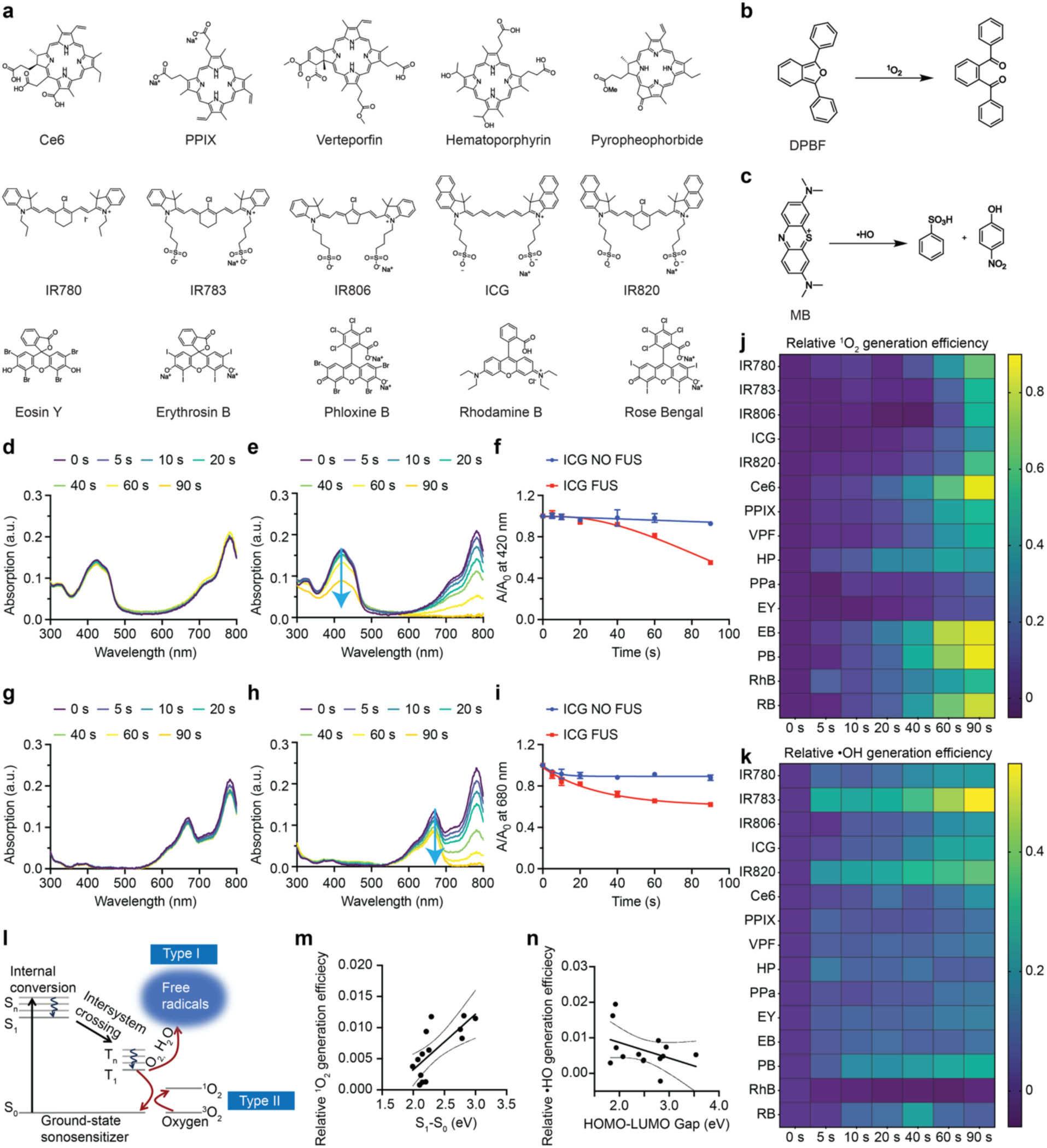
Screening of sonosensitizers for ultrasound-induced ROS generation and theoretical correlation modeling. (**a**) The small-molecule organic sonosensitizers investigated for ROS generation are categorized into three structural classes: porphyrin, cyanine, and xanthene derivatives. (**b**) Reaction scheme illustrating the detection of ^1^O_2_ using DPBF, whose characteristic absorption peak at 420 nm decreases proportionally with increasing ^1^O_2_ concentration. (**c**) Reaction mechanism of •OH detection using MB, whose 680 nm absorption peak decreases upon reaction with •OH. (**d**) UV-Vis spectra showing no significant change in DPBF absorption in the presence of ICG without ultrasound stimulation (1.5 MHz, 1.5 MPa, pulse 500 ms on, 500 ms off). (**e**) Time-dependent UV-Vis spectra demonstrating DPBF decomposition and ^1^O_2_ generation by ICG under ultrasound stimulation. (**f**) Quantitative analysis of DPBF decomposition induced by ultrasound in the presence of ICG compared to controls (n > 3 per group). (**g**) UV-Vis spectra indicating negligible MB decomposition in the absence of ultrasound. (**h**) Time-dependent UV-Vis spectra demonstrating MB degradation by •OH produced from ICG under ultrasound stimulation. (**i**) Quantitative analysis of MB decomposition with and without ultrasound irradiation in the presence of ICG (n > 3 per group). (**j**) Comparative analysis of ^1^O_2_ generation efficiency for fifteen sonosensitizers under identical ultrasound parameters. (**k**) Comparative analysis of •OH generation efficiency for fifteen sonosensitizers under identical ultrasound parameters. (**l**) Schematic Jablonski diagram illustrating ROS generation mechanisms of sonosensitizers activated by ultrasound. (**m**) Correlation plot demonstrating the positive relationship between relative ^1^O_2_ generation efficiency and S_1_-S_0_ energy gap. (**n**) Correlation plot showing the negative relationship between relative •OH generation efficiency and HOMO-LUMO energy gap.

### Sonosensitizer-optimized Mechanoluminescent Nanoparticles for Neuromodulation

The palette of organic mechanoluminescent nanoparticles developed herein covers a broad luminescence wavelength range from 459 nm to 592 nm, theoretically enabling activation or inhibition of most rhodopsins utilized in optogenetics.^47,48^ To validate the capability of these nanoparticles, we selected ChR2, eOPN3, and ChRmine as representative rhodopsins corresponding to blue, green, and red ML emissions, respectively. We systematically screened Sono-ML intensities of each nanoparticle type combined with fifteen different sonosensitizers. For blue ML, L012 NPs loaded with the sonosensitizer pyropheophorbide-a methyl ester (PPa) exhibited the highest intensity, significantly surpassing other combinations. Xanthene-based sonosensitizers generally demonstrate moderate luminescence intensities (**Figure 4a**). For green Sono-ML intended for eOPN3 inhibition, L012-FL NPs with PPa yielded the highest luminescence intensity; IR780 and EY produced ∼50% of that level (**Figure 4b**). For red ML applicable to ChRmine activation, L012-RhB NPs with EY displayed the brightest emission, while IR780, EB, PB, RhB, and RB also generated substantial red luminescence (**Figure 4c**). Based on these screens, optimized nanoparticle-sonosensitizer pairs were selected for subsequent sono-optogenetic experiments: L012 NPs with PPa for ChR2 (= 2.3-fold higher luminescence than the commonly used IR780 condition), L012-FL NPs with PPa for eOPN3 (= 1.8-fold), and L012-RhB NPs with EY for ChRmine (= 1.7-fold) (**Figure 4d–f**). Across formulations, blue-band ML intensity, which was not involved in further FRET during comparison of sonosensitizers, correlated strongly with relative ^1^O₂ generation efficiency (Pearson correlation r = 0.69; **Figure 4g**), underscoring ROS production as a key determinant of ML performance.

**Figure 4.**
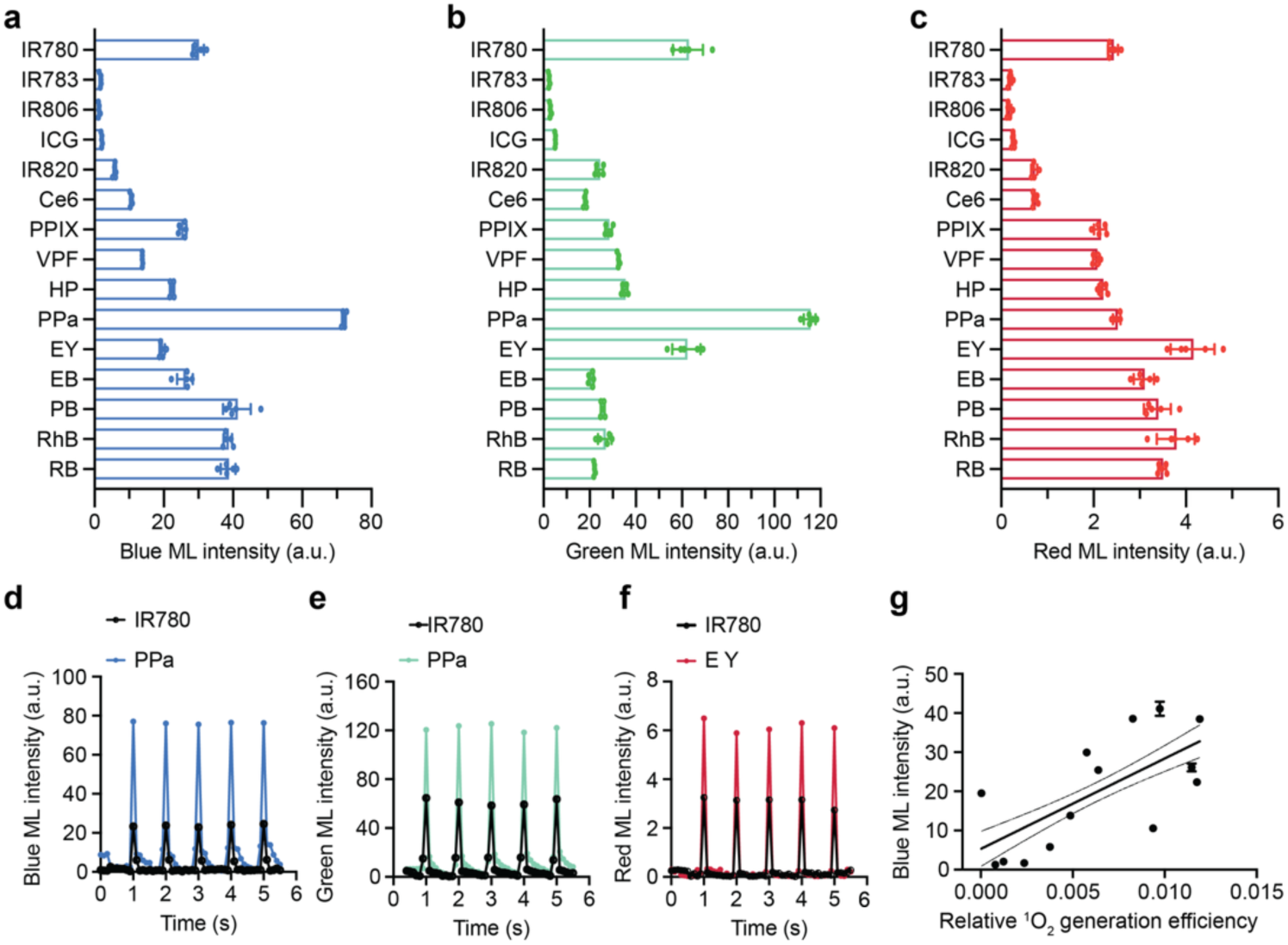
Comparative Sono-ML screening of fifteen sonosensitizers. (**a**) Comparative blue Sono-ML intensities of L012 NPs paired with fifteen sonosensitizers under FUS stimulation (1.5 MHz, 1.55 MPa, pulse 100 ms on, 900 ms off). (**b**) Comparative green Sono-ML intensities of L012-FL NPs combined with fifteen sonosensitizers under identical ultrasound conditions. (**c**) Comparative red Sono-ML intensities of L012-RhB NPs in combination with fifteen sonosensitizers under the same ultrasound parameters. (**d**) Representative Sono-ML intensity dynamics of L012 NPs combined with IR780 or PPa sonosensitizers under ultrasound stimulation (1.5 MHz, 1.55 MPa, pulse 100 ms on, 900 ms off). (**e**) Representative Sono-ML intensity dynamics of L012-FL NPs combined with IR780 or PPa sonosensitizers under the same ultrasound parameters. (**f**) Representative Sono-ML intensity dynamics of L012-RhB NPs combined with IR780 or EY sonosensitizers under identical ultrasound stimulation conditions. (**g**) Positive correlation analysis illustrating the relationship between relative ^1^O_2_ generation efficiency and blue Sono-ML intensity.

Next, we also investigate the potential of these mechanoluminescent nanoparticles in activating or inhibiting rhodopsin-expressing neurons under FUS stimulation. The primary neurons are transduced through adeno-associated viruses (AAVs) carrying rhodopsin genes (ChR2, eOPN3, and ChRmine), respectively. Neuronal activity was monitored through a microelectrode array (MEA) system. The experimental configuration involved placing a vial containing mechanoluminescent nanoparticles above the MEA well, with a miniaturized ultrasound transducer inserted to stimulate nanoparticles and facilitate photon emission, as depicted in **Figure 5a**. Initial experiments assessed ChR2 activation using ultrasound-triggered luminescence from L012 NPs. L012 showed a peak ML emission at ∼459 nm overlapping the ChR2 absorption maximum (**Figure 5b**).^49^ The stimulation sequence was 10 s baseline, 30 s ultrasound, and 10 s post-monitoring (**Figure 5b**). Real-time recordings showed increased spiking only under concurrent FUS and L012 NPs, with spike probabilities of approximately 97%, whereas controls lacking either component showed no ultrasound-dependent activation (**Figure 5c-d**). For inhibition, primary neurons expressing the green-sensitive rhodopsin eOPN3 were paired with L012-FL NPs.^50^ The green ML emission substantially overlapped the eOPN3 absorption spectrum (**Figure 5e**). The protocol used a 10 s baseline and 30 s continuous ultrasound-evoked green ML, with 1 Hz electrical drive from 10–40 s. Controls fired robustly during electrical stimulation, whereas simultaneous ultrasound and green ML reduced spike probability to approximately 17% (**Figure 5f-g**). For ChRmine, a red-shifted channelrhodopsin activated near 600 nm, L012-RhB NPs provided spectrally matched red ML (**Figure 5h**).^51^ Neurons expressing ChRmine exhibited synchronized spiking with approximately 84% spike probability only when both ultrasound and L012-RhB NPs were present; controls showed no activation (**Figure 5i-j**). Together, these results demonstrate bidirectional control of neuronal activity via spectrally matched mechanoluminescence under a standardized ultrasound regimen.

**Figure 5.**
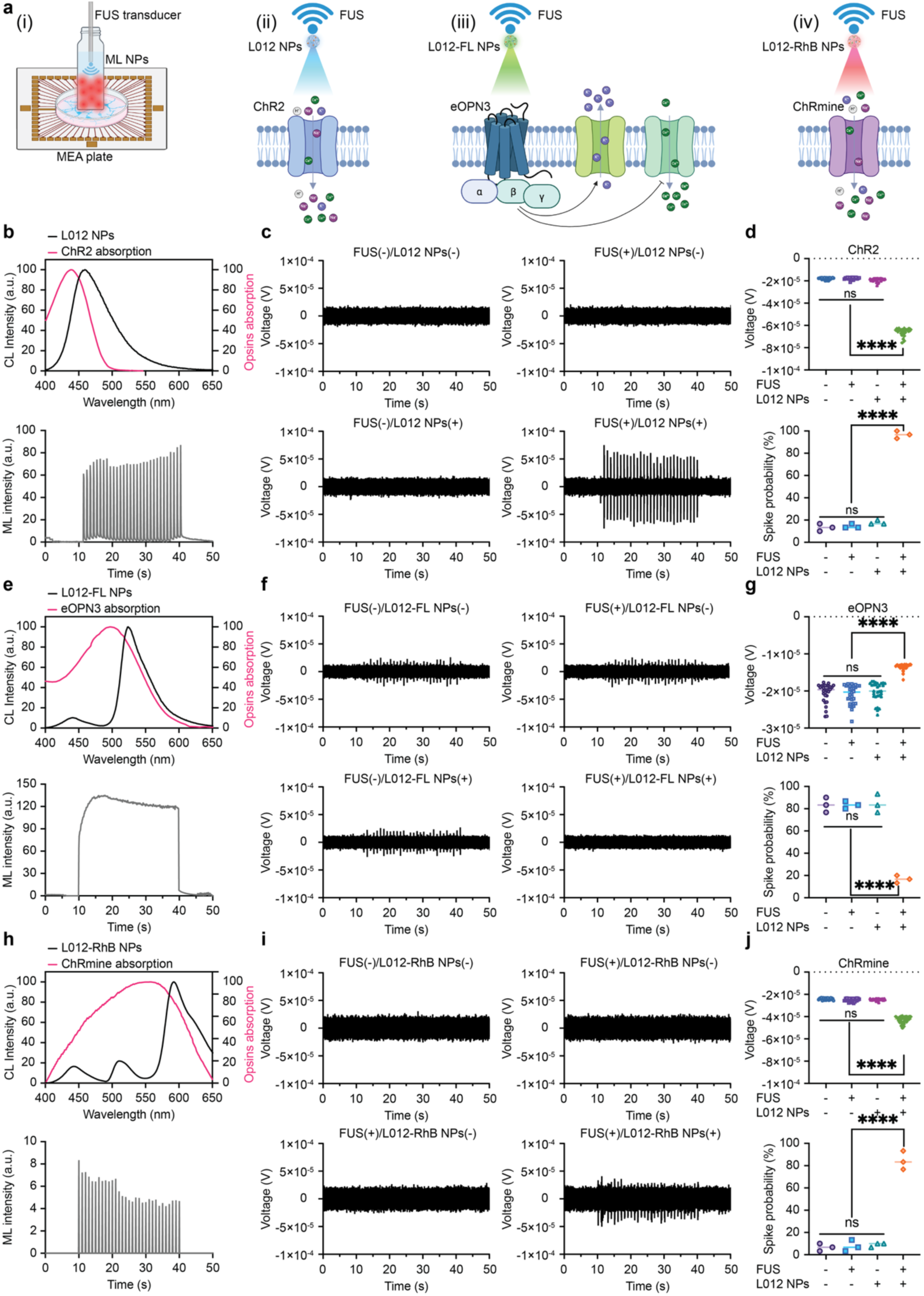
In vitro sono-optogenetics using mechanoluminescent nanoparticles in primary neurons expressing ChR2, eOPN3, and ChRmine. (**a**) Schematic illustration of the sono-optogenetic setup on a MEA system, showing: (i) experimental overview; (ii) ChR2 activation by L012 NPs under FUS; (iii) eOPN3 inhibition by L012-FL NPs; and (iv) ChRmine activation by L012-RhB NPs, created via BioRender.com. (**b**) Overlaid spectra of the CL emission from L012 NPs and the absorption profile of ChR2, demonstrating spectral compatibility. Below: representative Sono-ML trace used in MEA testing. (**c**) MEA recordings of ChR2-expressing neurons under four experimental conditions: FUS(–)/L012 NPs(–); FUS(+)/L012 NPs(–); FUS(–)/L012 NPs(+) and FUS(+)/L012 NPs(+). All recordings were conducted under standardized ultrasound conditions (1.5 MHz, 1.55 MPa, 100 ms on/900 ms off; stimulation from 10 s to 40 s). (**d**) Quantitative analysis of MEA signal changes across conditions and spike probability of ChR2-expressing neurons (n = 3 per group, two-way ANOVA and multiple comparisons test). (**e**) Overlaid spectra of L012-FL NPs and eOPN3 opsin absorption, confirming spectral overlap. Below: Sono-ML pattern used for the MEA experiment. (**f**) MEA recordings of eOPN3-expressing neurons under continuous electrical stimulation across four experimental conditions: FUS(–)/L012-FL NPs(–), FUS(+)/L012-FL NPs(–), FUS(–)/L012-FL NPs(+), and FUS(+)/L012-FL NPs(+). In the FUS(+)/L012-FL NPs(+) group, continuous Sono-ML irradiation was applied from 10 to 40 seconds. (**g**) Statistical analysis of MEA signal changes and spike probability of eOPN3-expressing neurons (n = 3 per group, two-way ANOVA and multiple comparisons test). (**h**) Overlaid spectra of L012-RhB NPs and ChRmine opsin absorption. Below: corresponding Sono-ML trace used in the MEA test. (**i**) MEA recordings of ChRmine-expressing neurons under the same four experimental conditions and ultrasound parameters as in (**c**). (**j**) Quantitative analysis of MEA signal variations and spike probability of ChRmine-expressing neurons across all tested groups (n = 3 per group, two-way ANOVA and multiple comparisons test). All plots show mean ± SEM unless otherwise mentioned. *P < 0.05, **P < 0.01, ***P < 0.001, ****P < 0.0001; ns, not significant.

## Conclusion

In summary, we developed a wavelength-tunable palette of organic mechanoluminescent nanoparticles capable of multicolor photon emission upon focused ultrasound stimulation. Through the stepwise optimization of chemiluminescence and systematic evaluation of mechanoluminescence behavior, we achieved ultrasound-triggered emission tunability ranging from 459 nm to 592 nm. Furthermore, fifteen distinct small molecule sonosensitizers were comprehensively analyzed to elucidate the mechanistic relationships between energy levels, ROS generation efficiency, and subsequent light emission. These insights enabled the rational selection of nanoparticle–sonosensitizer combinations tailored for activation of both excitatory and inhibitory opsins with spectral sensitivities spanning the visible range. This study demonstrates a robust and versatile platform for non-invasive, multicolor, and opsin-targeted neuromodulation, expanding the toolbox that could be used for sono-optogenetics.

## Experimental Section/Methods

The experimental details and all characterization are listed in the Supporting Information. All procedures were designed according to the National Institute of Health Guide for the Care and Use of Laboratory Animals, were approved by the Institutional Animal Care and Use committee at the University of Texas at Austin and were supported via the Animal Resources Center at the University of Texas at Austin.

## Supporting information

The Supporting Information includes detailed fabrication methods for the mechanoluminescent nanoparticles and quantitative assessments of reactive oxygen species (ROS) generation efficiencies for the 15 evaluated sonosensitizers. This material is freely accessible online at http://pubs.acs.org.

## Acknowledgement

Dr. Huiliang Wang acknowledges funding support from NIH Maximizing Investigators’ Research Award (National Institute of General Medical Sciences 1R35GM147408), XiNational Science Foundation (NSF) CAREER award (2340964), UT Austin Portugal Exploratory Research Projects Grant, Robert A. Welch Foundation Grant, Texas Biologics Pilot Grant, and Craig H. Neilsen Foundation Pilot Research Grant.

## TOC Graphic

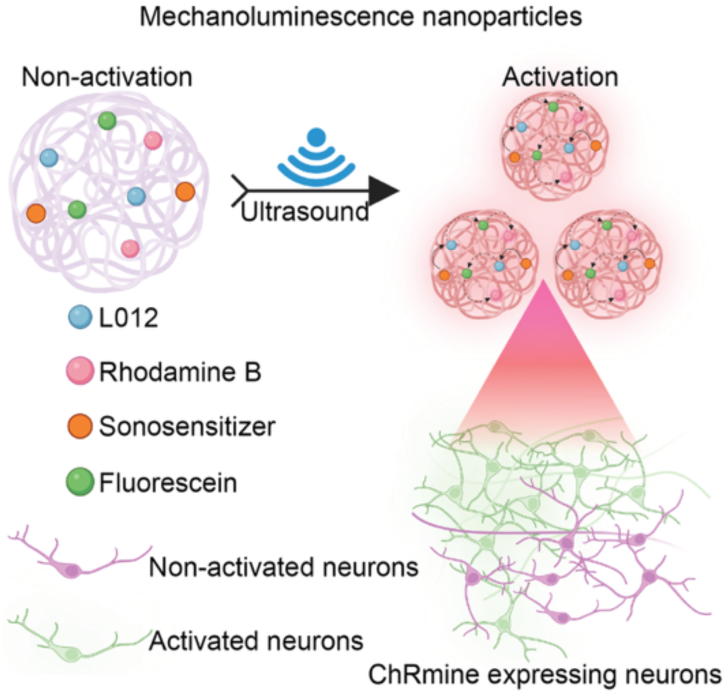

