## Supporting Information for "Multicolored, Sonosensitizer-optimized Organic Mechanoluminescent Nanoparticles for Functional Sono-optogenetics"

*Corresponding author

**Experimental Section/Methods**

**Materials**: β-Cyclodextrin, $CoCl_2\cdot 6H_2O$, L012, fluorescein sodium salt, eosin Y, phloxine B, rhodamine B, Ce6, protoporphyrin IX, verteporfin, hematoporphyrin, pyropheophorbide a methyl ester, IR780, IR783, IR806, ICG, IR820, erythrosin B, rose bengal, 1,3-diphenylisobenzofuran and methylene blue were ordered from Sigma-Aldrich. pAAV-hSyn-hChR2(H134R)-EYFP, pAAV-CaMKIIa(0.4)-eOPN3-mScarlet-WPRE and pAAV-hSyn-ChRmine-mScarlet-Kv2.1-WPRE were ordered from Addgene.

**Characterization**: The hydrodynamic diameter and surface potential of these nanoparticles were evaluated through dynamic light scattering (DLS, Zetasizer Nano-ZS from Malvern Instruments). The UV-Vis spectra were measured via Eppendorf Biospectrometer. The chemiluminescence spectra were evaluated through Fluorolog3 Fluorimeter. The focused ultrasound stimulation was derived from Image Guided Therapy. The optical images of these chemiluminescence nanoparticles were acquired through Fluorescence In Vivo Imaging System (FOBI). The microelectrode array (MEA) were measured via Maestro Edge from Axion Biosystems.

**Preparation of nanoparticles**: β-CD-assisted alkaline coordination strategy was used to formulate these nanoparticles.[1] First, chemical stock solutions are prepared based on the following description: 20 mg NaOH was dissolved into 50 mL deionized water (DI water) to prepare 10 mM NaOH stock solution; 1.12 mg L012 was dissolved into 0.6 mL 10 mM NaOH stock solution to prepare 6 mM L012 stock solution; 1.33 mg IR780 was dissolved into 1 mL mixed solvent (0.5 mL MeOH+0.5 mL $H_2O$) to prepare 2 mM IR780 stock solution; 265 mg β-CD was dissolved into 5 mL DI water to prepare 46.7 mM β-CD stock solution; 12.11 mg Tris base was dissolved into 1 mL DI water to prepare 100 mM Tris base stock solution; 4.76 mg $CoCl_2\cdot 6H_2O$ was dissolved into 1 mL DI water to prepare 20 mM $Co^{2+}$ stock solution; 3.76 mg fluorescein sodium salt was dissolved into 0.2 mL DI water to prepare 50 mM fluorescein stock solution; 6.92 mg eosin Y disodium salt was dissolved into 0.2 mL DI water to prepare 50 mM eosin Y stock solution; 8.30 mg phloxine B was dissolved into 0.2 mL DI water to prepare 50 mM phloxine B stock solution and 4.79 mg rhodamine B was dissolved into 0.2 mL DI water to prepare 50 mM rhodamine B stock solution. Secondly, to prepare L012 NPs, 110 μL L012 stock solution, 30 μL IR780 stock solution, 90 μL β-CD stock solution 37.2 μL Tris base stock solution, 0.6 μL $Co^{2+}$ stock solution and 32.2 μL DI water were mixed together and incubated at 37°C in shaking incubator for 30 minutes to get the L012 NPs. To prepare L012-FL NPs, 110 μL L012 stock solution, 30 μL IR780 stock solution, 90 μL β-CD stock solution 37.2 μL Tris base stock solution, 0.6 μL $Co^{2+}$ stock solution, 7.2 μL fluorescein stock solution and 25 μL DI water were mixed together and incubated at 37°C in shaking incubator for 30 minutes to get the L012-FL NPs. L012-EY NPs were prepared by adding 110 μL L012 stock solution, 30 μL IR780 stock solution, 90 μL β-CD stock solution, 37.2 μL Tris base stock solution, 0.6 μL $Co^{2+}$ stock solution, 7.2 μL fluorescein stock solution, 2.4

μL EY stock solution and 22.6 μL DI water together and incubated at 37°C in shaking incubator for 30 minutes. L012-PB/RhB NPs were prepared by adding 110 μL L012 stock solution, 30 μL IR780 stock solution, 90 μL β-CD stock solution, 37.2 μL Tris base stock solution, 0.6 μL $Co^{2+}$ stock solution, 7.2 μL fluorescein stock solution, 3.6 μL PB/RhB stock solution and 21.4 μL DI water together and incubated at 37°C in shaking incubator for 30 minutes. To prepare a series of wavelength-tunable chemiluminescence NPs, 30 μL IR780 stock solution was not added into the nanoparticle system and 30 μL $H_2O_2$ was added to initiate the chemiluminescence.

**Chemiluminescence spectra of a series of wavelength-tunable nanoparticles**:

A mixture of 0.3 mL wavelength-tunable chemiluminescent nanoparticle solution and 30 μL $H_2O_2$ was transferred to a fluorescence cuvette and placed in a spectrofluorometer (FluoroMax-4). Chemiluminescence spectra were recorded without excitation light. Absorption spectra for ChR2, eOPN3, and ChRmine were taken from prior work and digitized using WebPlotDigitizer.

**Study of optical images of these chemiluminescent nanoparticles**:

A mixture of 0.3 mL wavelength-tunable chemiluminescent nanoparticle solution and 30 μL $H_2O_2$ was dispensed into a black 96-well polystyrene microplate, which was then imaged on a FOBI system. The optical chemiluminescence images were acquired at 30 frames per second with gain 100.

**FUS triggered luminescence from a series of mechanoluminescence nanoparticles**:

A 1 mL aliquot of wavelength-tunable mechanoluminescent nanoparticle solution was added to a 2 mL glass vial and coupled to a 1.5 MHz FUS transducer (Image Guided Therapy). The gap between the transducer and vial was filled with ultrasound gel. A monochrome camera (Zelux CS165MU1/M, 1.6 MP; Thorlabs) recorded light emission in a dark room. We first measured emission at a 1 Hz repetition frequency (100 ms on, 900 ms off) while varying peak pressure from 0 to 1.67 MPa. We then evaluated emission at 1.55 MPa across pulse rates of 2 Hz (100 ms on, 400 ms off), 4 Hz (100 ms on, 150 ms off), and 8 Hz (100 ms on, 25 ms off). At 1.55 MPa and 1 Hz, we next varied pulse duration: 100 ms (100 ms on, 900 ms off), 300 ms (300 ms on, 700 ms off), 500 ms (500 ms on, 500 ms off), and 900 ms (900 ms on, 100 ms off). To assess effective ML duration, ultrasound was applied in 1 s on/1 s off cycles for 1200 s. All data were analyzed in ImageJ.

**Computational methods:**

Ground-state geometries ($S_0$) of all sonosensitizers were optimized with DFT using the B3LYP functional as implemented in ORCA. The def2-SVP basis set and RIJCOSX approximation with the corresponding def2/J auxiliary basis were applied. Vertical singlet excitation energies ($S_1$) were calculated by TD-DFT at the same level of theory, and HOMO, LUMO and HOMO - LUMO gaps were obtained from the Kohn-Sham orbital eigenvalues of the optimized $S_0$ structures. The $S_1$ - $S_0$ gaps were taken from the lowest singlet vertical excitation energies.

**Detection of the generation of $^1O_2$**:

1 mL sonosensitizer (0.1 mg/mL in the 10% DMSO aqueous solution) with 30 μL 1 mg/mL DPBF (dissolved in methanol) were mixed together. The UV-Vis characteristic absorption peak of DPBF at 420 nm was used to track the generation of $^1O_2$. The mixture was treated with or without FUS irradiation (1.5 MHz, 1.5 MPa) and extracted out 10 μL for UV-Vis spectrum tests. The absorbance change of DPBF at 420 nm was used to quantify the generation of $^1O_2$.

**Detection of the generation of $^{\bullet}OH$**:

1 mL sonosensitizer (0.1 mg/mL in the 10% DMSO aqueous solution) with 30 μL 1 mg/mL MB were mixed together. The UV-Vis characteristic absorption peak of MB at 680 nm was used to track the generation of $^{\bullet}OH$. The mixture was treated with or without FUS irradiation (1.5 MHz, 1.5 MPa) and extracted out 10 μL for UV-Vis spectrum tests. The absorbance change of MB at 680 nm was used to quantify the generation of $^{\bullet}OH$.

**MEA tests of in-vitro sono-optogenetics**:

We evaluated the sono-mechanoluminescence triggered firing ChR2, EOPN3 and ChRmine expressing primary neurons. Primary cortical neurons were used in our tests. Briefly, the pregnant C57BL/6 mouse (20–26 g; 8 weeks old; Jackson Laboratory) was sacrificed when the pups were 15.5 days old, and these pups' brains were used to prepare the primary cortical neurons. The 24 MEA well plates were coated with poly-l-ornithine (0.2 mg/mL) at 37 °C for 2 h and washed with PBS several times to remove excessive poly-l-ornithine, then warmed at 37 °C cell incubator before use. The dissociated cortical neurons were plated into the plate with suitable cell density, and cultured in neurobasal medium with 10 % B27, glutamine, penicillin, and streptomycin. After incubating for 2 days at 37 °C under 7% CO2, the glial inhibitor 5-fluoro-2′- deoxyuridine (0.1 mM) was added. After 4 days of incubation, 0.5 μL pAAV-hSyn-hChR2(H134R)-EYFP, pAAV-CaMKIIa(0.4)-eOPN3-mScarlet-WPRE and pAAV-hSyn-ChRmine-mScarlet-Kv2.1-WPRE were added to infect the neurons in different wells. After another 7 days of incubation, the rhodopsins ChR2, EOPN3 and ChRmine were successfully expressed in the neurons for MEA test. The vail filling with 1 mL wavelength-tunable mechanoluminescence nanoparticle solution was fixed over the cells, and the FUS irradiation (1.55MPa, pulse 100 ms on 900 ms off) was given to activate the system for light generation. For the inhibitory rhodopsins eOPN3, we initiate the electrical stimulation ( pulse 100 ms on 900 ms off) for all the four groups. The MEA signals were collected and recorded via Maestro Edge.

**Figures**

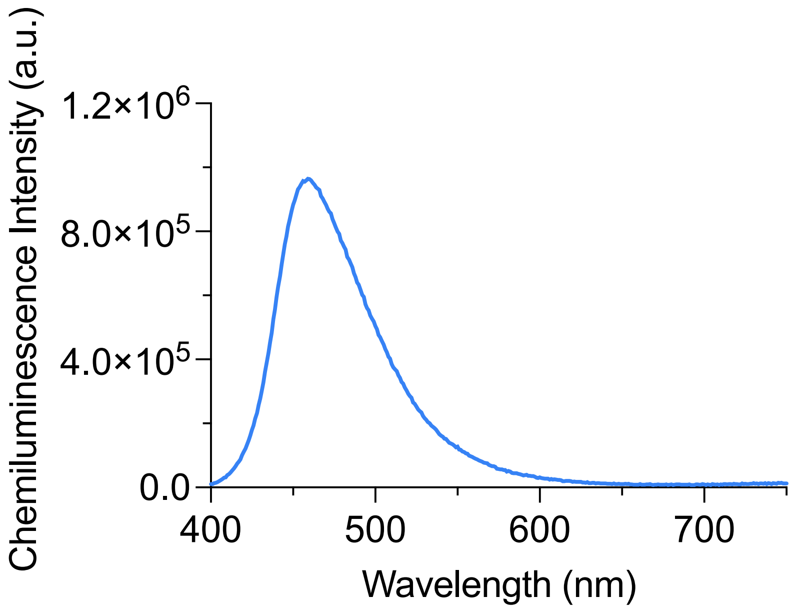


**Figure S1**. The chemiluminescence spectrum of L012 NPs. The emission peak of L012 NPs is at around 459 nm.

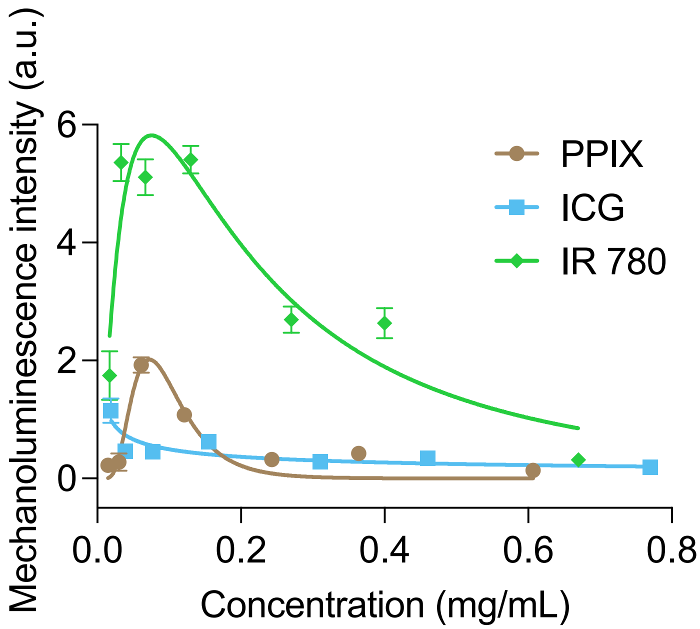


**Figure S2**. The sonosensitizer concentration dependent mechanoluminescence intensity of L012 NPs. The highest mechanoluminescence intensity is at around 0.1 mg/mL.

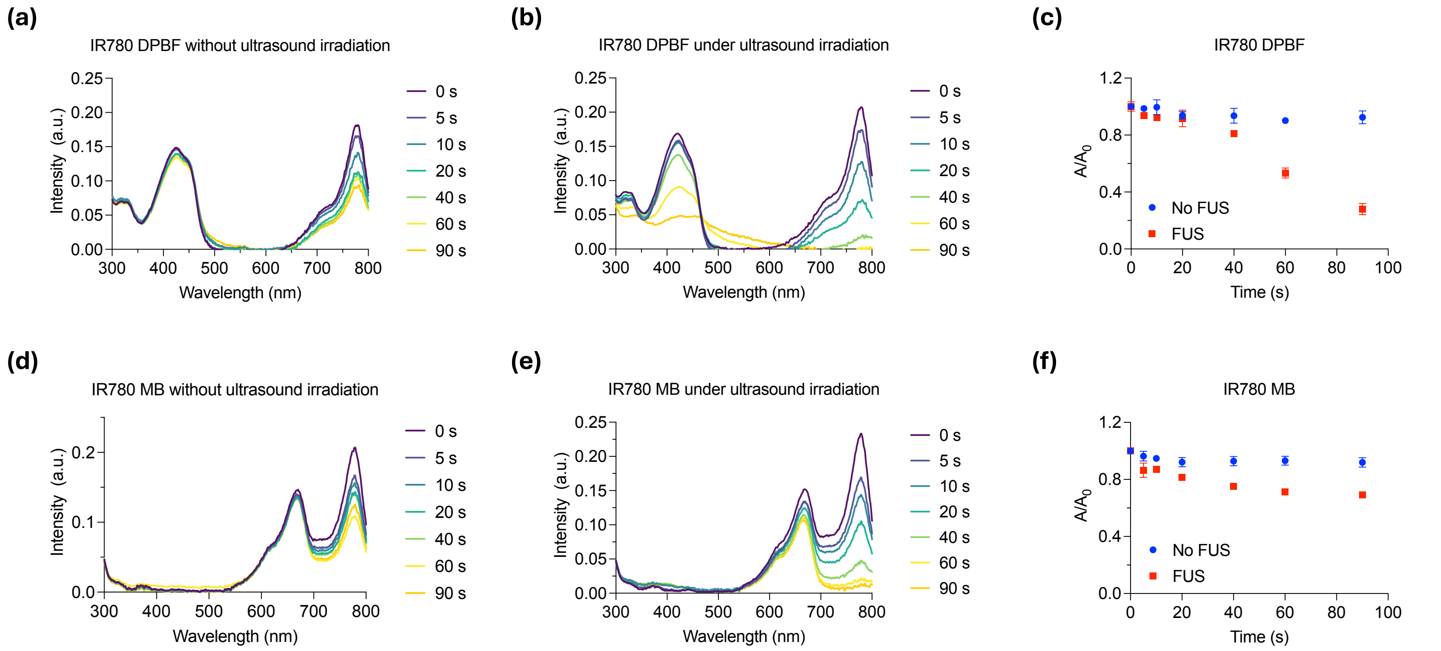


**Figure S3**. The ultrasound-induced ROS generation of sonosensitizer IR780. (a) UV-Vis spectra showing no significant change in DPBF absorption in the presence of IR780 without ultrasound stimulation (1.5 MHz, 1.5 MPa, pulse 500 ms on, 500 ms off). (b) Time-dependent UV-Vis spectra demonstrating DPBF decomposition and $^1O_2$ generation by IR780 under ultrasound stimulation. (c) Quantitative analysis of DPBF decomposition induced by ultrasound in the presence of IR780 compared to controls (n > 3 per group). (d) UV-Vis spectra indicating negligible MB decomposition in the absence of ultrasound. (e) Time-dependent UV-Vis spectra demonstrating MB degradation by •OH produced from IR780 under ultrasound stimulation. (f) Quantitative analysis of MB decomposition with and without ultrasound irradiation in the presence of IR780 (n > 3 per group). All plots show mean ± SEM unless otherwise mentioned.

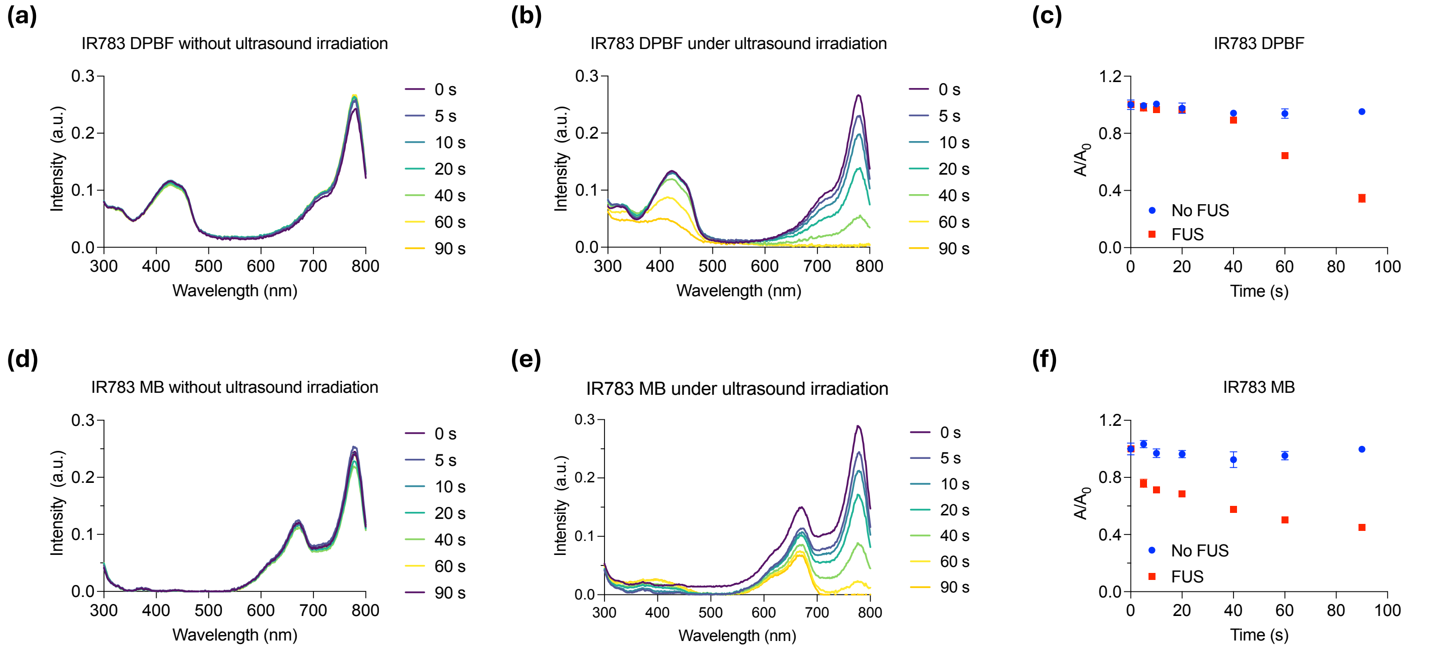


**Figure S4**. The ultrasound-induced ROS generation of sonosensitizer IR783. (a) UV-Vis spectra showing no significant change in DPBF absorption in the presence of IR783 without ultrasound stimulation (1.5 MHz, 1.5 MPa, pulse 500 ms on, 500 ms off). (b) Time-dependent UV-Vis spectra demonstrating DPBF decomposition and $^1O_2$ generation by IR783 under ultrasound stimulation. (c) Quantitative analysis of DPBF decomposition induced by ultrasound in the presence of IR783 compared to controls (n > 3 per group). (d) UV-Vis spectra indicating negligible MB decomposition in the absence of ultrasound. (e) Time-dependent UV-Vis spectra demonstrating MB degradation by •OH produced from IR783 under ultrasound stimulation. (f) Quantitative analysis of MB decomposition with and without ultrasound irradiation in the presence of IR783 (n > 3 per group). All plots show mean ± SEM unless otherwise mentioned.

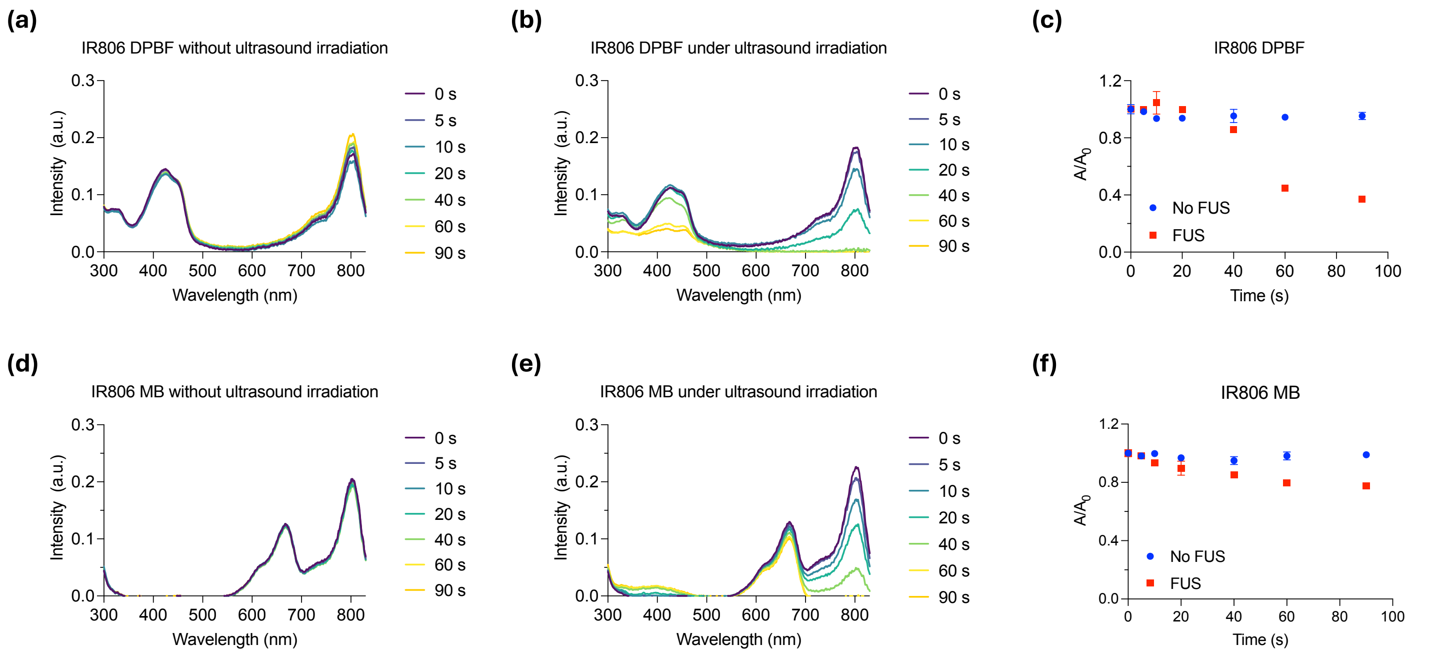


**Figure S5**. The ultrasound-induced ROS generation of sonosensitizer IR806. (a) UV-Vis spectra showing no significant change in DPBF absorption in the presence of IR806 without ultrasound stimulation (1.5 MHz, 1.5 MPa, pulse 500 ms on, 500 ms off). (b) Time-dependent UV-Vis spectra demonstrating DPBF decomposition and $^1O_2$ generation by IR806 under ultrasound stimulation. (c) Quantitative analysis of DPBF decomposition induced by ultrasound in the presence of IR806 compared to controls ($n > 3$ per group). (d) UV-Vis spectra indicating negligible MB decomposition in the absence of ultrasound. (e) Time-dependent UV-Vis spectra demonstrating MB degradation by •OH produced from IR806 under ultrasound stimulation. (f) Quantitative analysis of MB decomposition with and without ultrasound irradiation in the presence of IR806 ($n > 3$ per group). All plots show mean ± SEM unless otherwise mentioned.

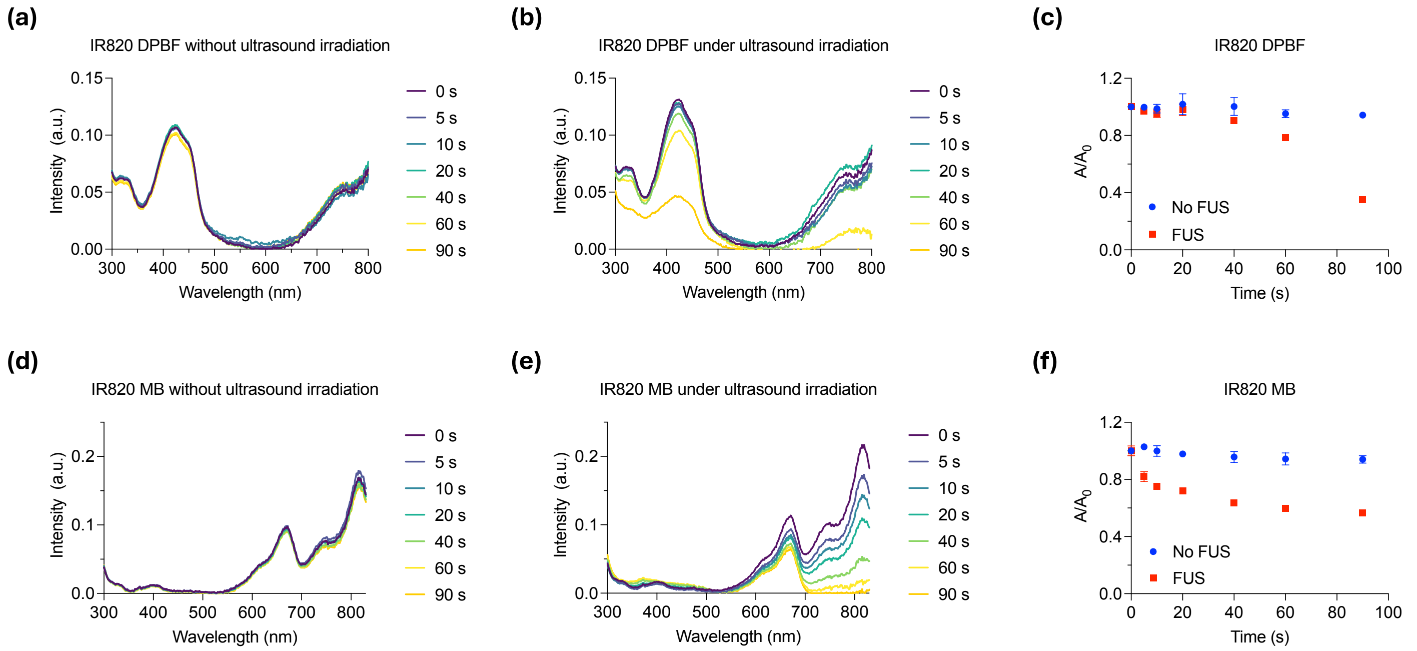


**Figure S6**. The ultrasound-induced ROS generation of sonosensitizer IR820. (a) UV-Vis spectra showing no significant change in DPBF absorption in the presence of IR820 without ultrasound stimulation (1.5 MHz, 1.5 MPa, pulse 500 ms on, 500 ms off). (b) Time-dependent UV-Vis spectra demonstrating DPBF decomposition and $^1O_2$ generation by IR820 under ultrasound stimulation. (c) Quantitative analysis of DPBF decomposition induced by ultrasound in the presence of IR820 compared to controls ($n > 3$ per group). (d) UV-Vis spectra indicating negligible MB decomposition in the absence of ultrasound. (e) Time-dependent UV-Vis spectra demonstrating MB degradation by •OH produced from IR820 under ultrasound stimulation. (f) Quantitative analysis of MB decomposition with and without ultrasound irradiation in the presence of IR820 ($n > 3$ per group). All plots show mean ± SEM unless otherwise mentioned.

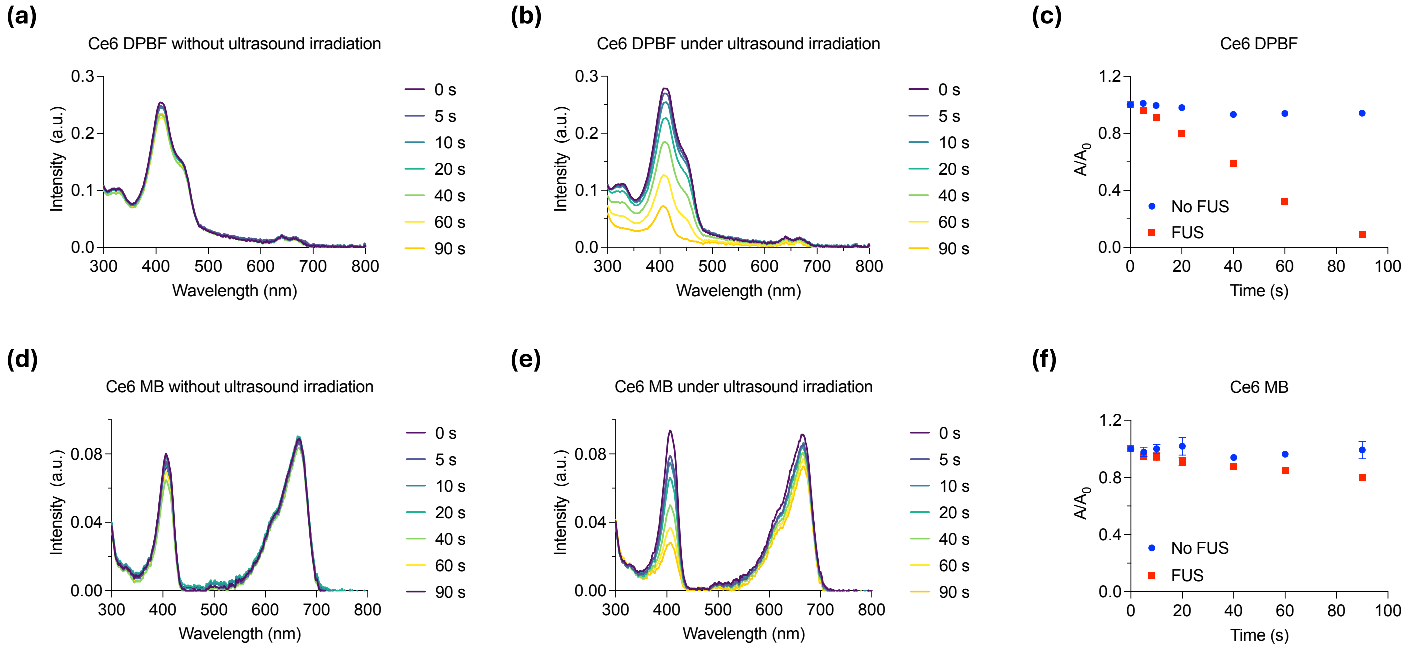


**Figure S7**. The ultrasound-induced ROS generation of sonosensitizer Ce6. (a) UV-Vis spectra showing no significant change in DPBF absorption in the presence of Ce6 without ultrasound stimulation (1.5 MHz, 1.5 MPa, pulse 500 ms on, 500 ms off). (b) Time-dependent UV-Vis spectra demonstrating DPBF decomposition and $^1O_2$ generation by Ce6 under ultrasound stimulation. (c) Quantitative analysis of DPBF decomposition induced by ultrasound in the presence of Ce6 compared to controls ($n > 3$ per group). (d) UV-Vis spectra indicating negligible MB decomposition in the absence of ultrasound. (e) Time-dependent UV-Vis spectra demonstrating MB degradation by •OH produced from Ce6 under ultrasound stimulation. (f) Quantitative analysis of MB decomposition with and without ultrasound irradiation in the presence of Ce6 ($n > 3$ per group). All plots show mean ± SEM unless otherwise mentioned.

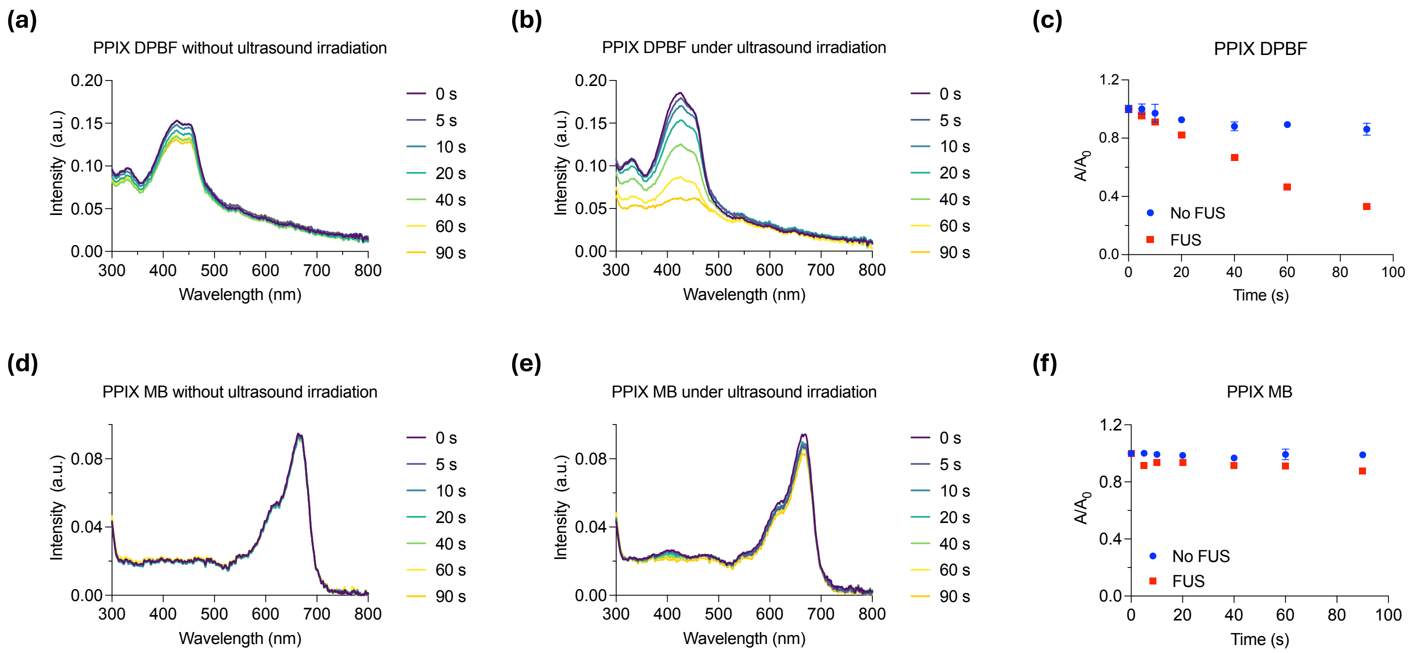


**Figure S8**. The ultrasound-induced ROS generation of sonosensitizer PPIX. (a) UV-Vis spectra showing no significant change in DPBF absorption in the presence of PPIX without ultrasound stimulation (1.5 MHz, 1.5 MPa, pulse 500 ms on, 500 ms off). (b) Time-dependent UV-Vis spectra demonstrating DPBF decomposition and $^1O_2$ generation by PPIX under ultrasound stimulation. (c) Quantitative analysis of DPBF decomposition induced by ultrasound in the presence of PPIX compared to controls ($n > 3$ per group). (d) UV-Vis spectra indicating negligible MB decomposition in the absence of ultrasound. (e) Time-dependent UV-Vis spectra demonstrating MB degradation by •OH produced from PPIX under ultrasound stimulation. (f) Quantitative analysis of MB decomposition with and without ultrasound irradiation in the presence of PPIX ($n > 3$ per group). All plots show mean ± SEM unless otherwise mentioned.

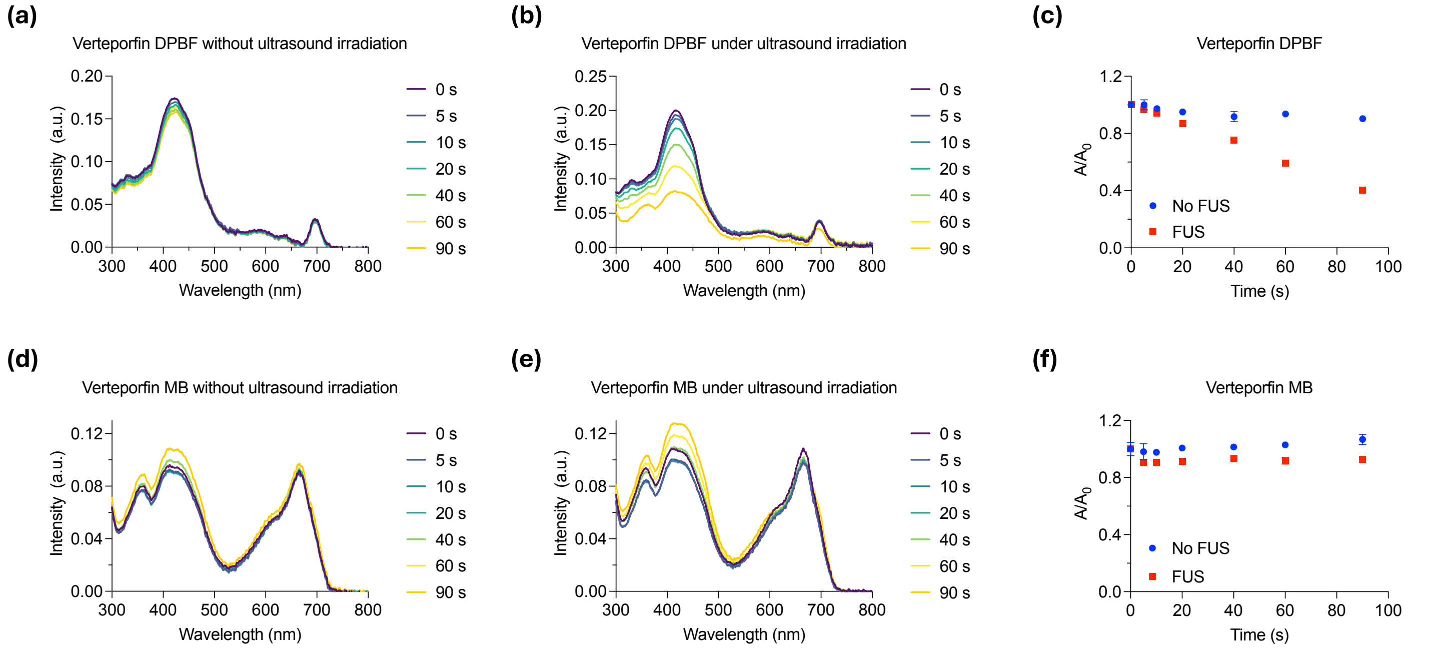


**Figure S9**. The ultrasound-induced ROS generation of sonosensitizer Verteporfin. (a) UV-Vis spectra showing no significant change in DPBF absorption in the presence of verteporfin without ultrasound stimulation (1.5 MHz, 1.5 MPa, pulse 500 ms on, 500 ms off). (b) Time-dependent UV-Vis spectra demonstrating DPBF decomposition and $^1O_2$ generation by verteporfin under ultrasound stimulation. (c) Quantitative analysis of DPBF decomposition induced by ultrasound in the presence of verteporfin compared to controls ($n > 3$ per group). (d) UV-Vis spectra indicating negligible MB decomposition in the absence of ultrasound. (e) Time-dependent UV-Vis spectra demonstrating MB degradation by •OH produced from verteporfin under ultrasound stimulation. (f) Quantitative analysis of MB decomposition with and without ultrasound irradiation in the presence of verteporfin ($n > 3$ per group). All plots show mean ± SEM unless otherwise mentioned.

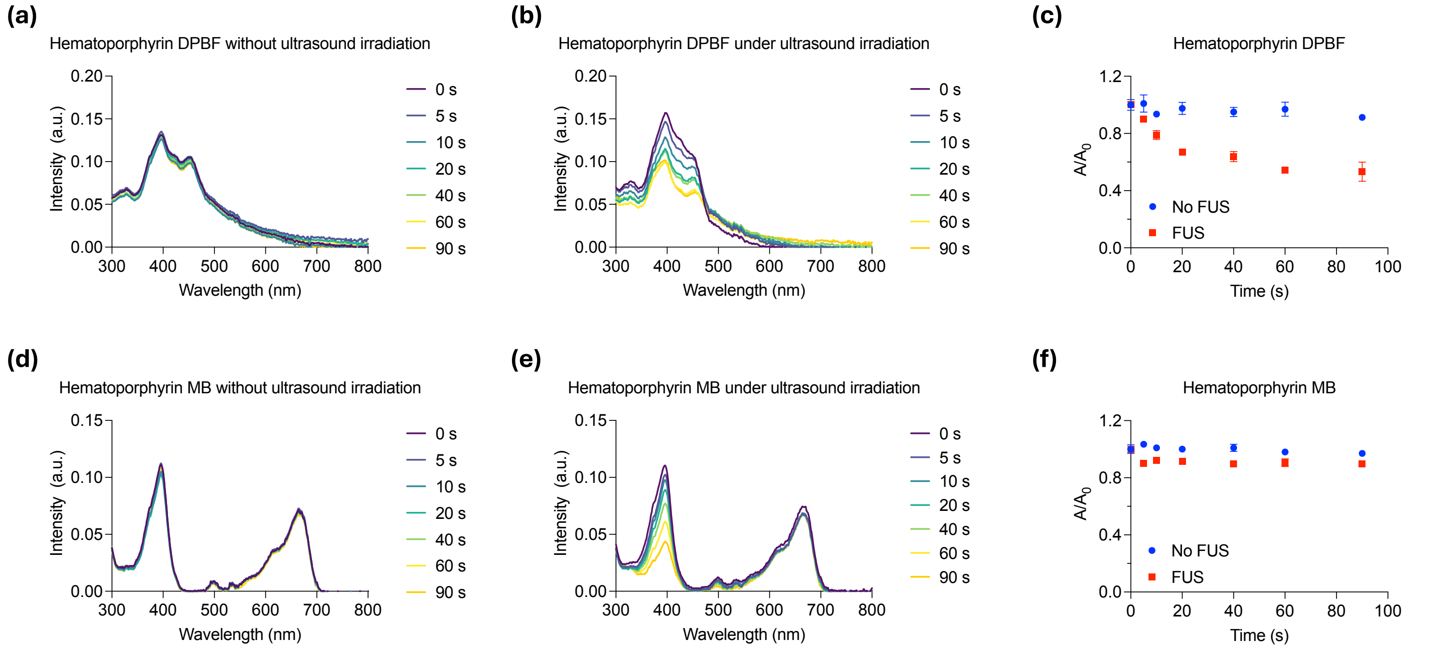


**Figure S10**. The ultrasound-induced ROS generation of sonosensitizer Hematoporphyrin. (a) UV-Vis spectra showing no significant change in DPBF absorption in the presence of hematoporphyrin without ultrasound stimulation (1.5 MHz, 1.5 MPa, pulse 500 ms on, 500 ms off). (b) Time-dependent UV-Vis spectra demonstrating DPBF decomposition and $^1O_2$ generation by hematoporphyrin under ultrasound stimulation. (c) Quantitative analysis of DPBF decomposition induced by ultrasound in the presence of hematoporphyrin compared to controls (n > 3 per group). (d) UV-Vis spectra indicating negligible MB decomposition in the absence of ultrasound. (e) Time-dependent UV-Vis spectra demonstrating MB degradation by •OH produced from hematoporphyrin under ultrasound stimulation. (f) Quantitative analysis of MB decomposition with and without ultrasound irradiation in the presence of hematoporphyrin (n > 3 per group). All plots show mean ± SEM unless otherwise mentioned.

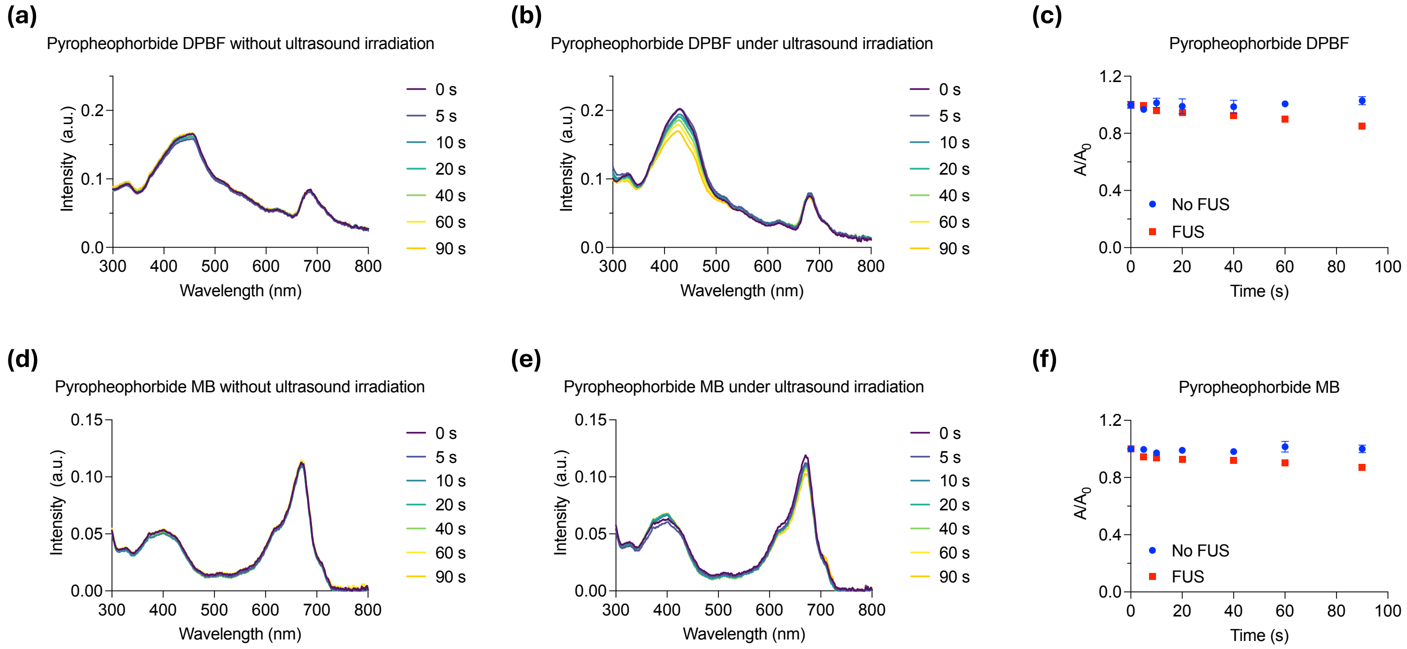


**Figure S11**. The ultrasound-induced ROS generation of sonosensitizer Pyropheophorbide a methyl ester. (a) UV-Vis spectra showing no significant change in DPBF absorption in the presence of pyropheophorbide a methyl ester without ultrasound stimulation (1.5 MHz, 1.5 MPa, pulse 500 ms on, 500 ms off). (b) Time-dependent UV-Vis spectra demonstrating DPBF decomposition and $^1O_2$ generation by pyropheophorbide a methyl ester under ultrasound stimulation. (c) Quantitative analysis of DPBF decomposition induced by ultrasound in the presence of pyropheophorbide a methyl ester compared to controls (n > 3 per group). (d) UV-Vis spectra indicating negligible MB decomposition in the absence of ultrasound. (e) Time-dependent UV-Vis spectra demonstrating MB degradation by •OH produced from pyropheophorbide a methyl ester under ultrasound stimulation. (f) Quantitative analysis of MB decomposition with and without ultrasound irradiation in the presence of pyropheophorbide a methyl ester (n > 3 per group). All plots show mean ± SEM unless otherwise mentioned.

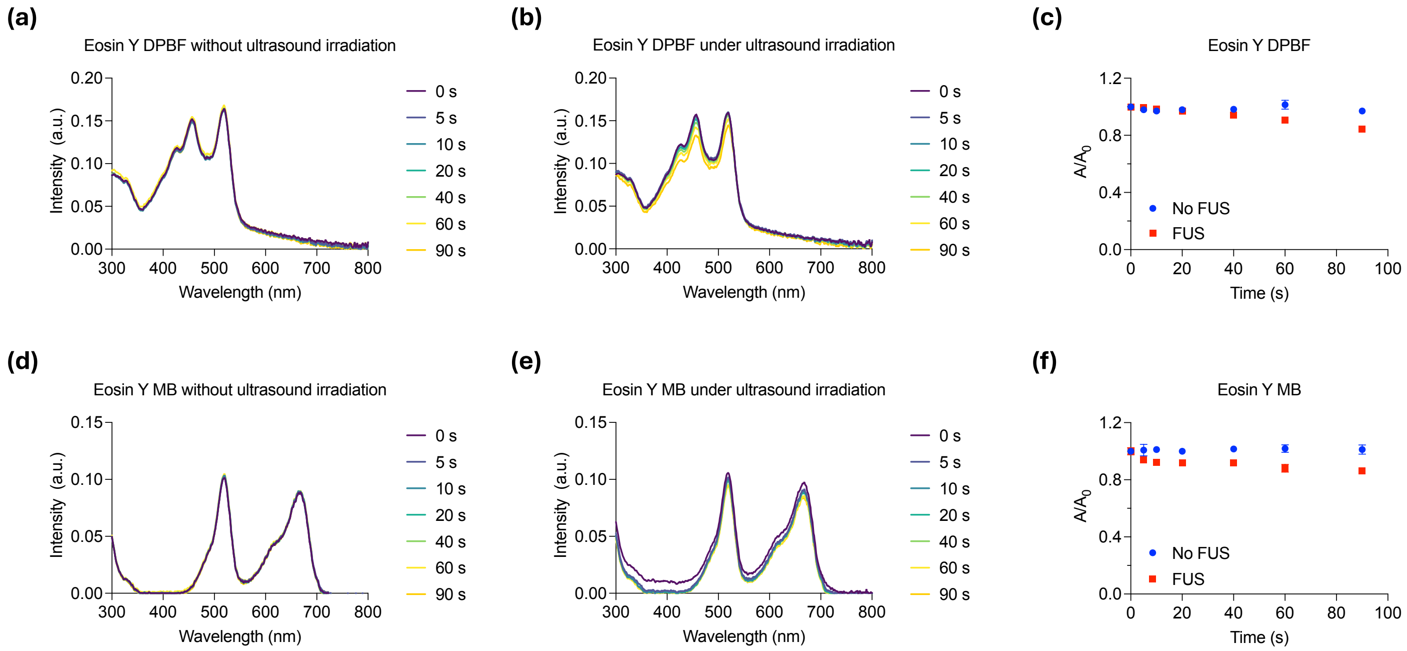


**Figure S12**. The ultrasound-induced ROS generation of sonosensitizer Eosin Y. (a) UV-Vis spectra showing no significant change in DPBF absorption in the presence of Eosin Y without ultrasound stimulation (1.5 MHz, 1.5 MPa, pulse 500 ms on, 500 ms off). (b) Time-dependent UV-Vis spectra demonstrating DPBF decomposition and $^1O_2$ generation by Eosin Y under ultrasound stimulation. (c) Quantitative analysis of DPBF decomposition induced by ultrasound in the presence of Eosin Y compared to controls (n > 3 per group). (d) UV-Vis spectra indicating negligible MB decomposition in the absence of ultrasound. (e) Time-dependent UV-Vis spectra demonstrating MB degradation by •OH produced from Eosin Y ultrasound stimulation. (f) Quantitative analysis of MB decomposition with and without ultrasound irradiation in the presence of Eosin Y (n > 3 per group). All plots show mean ± SEM unless otherwise mentioned.

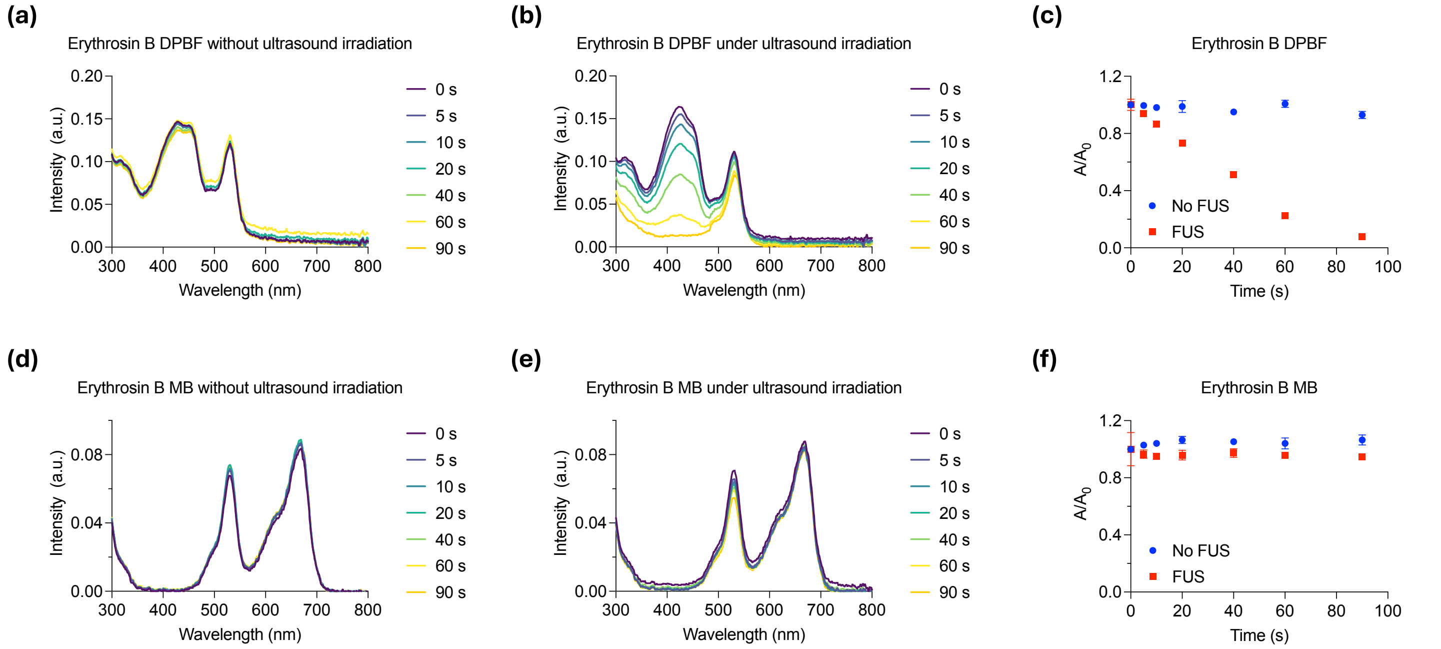


**Figure S13**. The ultrasound-induced ROS generation of sonosensitizer Erythrosin B. (a) UV-Vis spectra showing no significant change in DPBF absorption in the presence of Erythrosin B without ultrasound stimulation (1.5 MHz, 1.5 MPa, pulse 500 ms on, 500 ms off). (b) Time-dependent UV-Vis spectra demonstrating DPBF decomposition and $^1O_2$ generation by Erythrosin B under ultrasound stimulation. (c) Quantitative analysis of DPBF decomposition induced by ultrasound in the presence of Erythrosin B compared to controls ($n > 3$ per group). (d) UV-Vis spectra indicating negligible MB decomposition in the absence of ultrasound. (e) Time-dependent UV-Vis spectra demonstrating MB degradation by •OH produced from Erythrosin B ultrasound stimulation. (f) Quantitative analysis of MB decomposition with and without ultrasound irradiation in the presence of Erythrosin B ($n > 3$ per group). All plots show mean ± SEM unless otherwise mentioned.

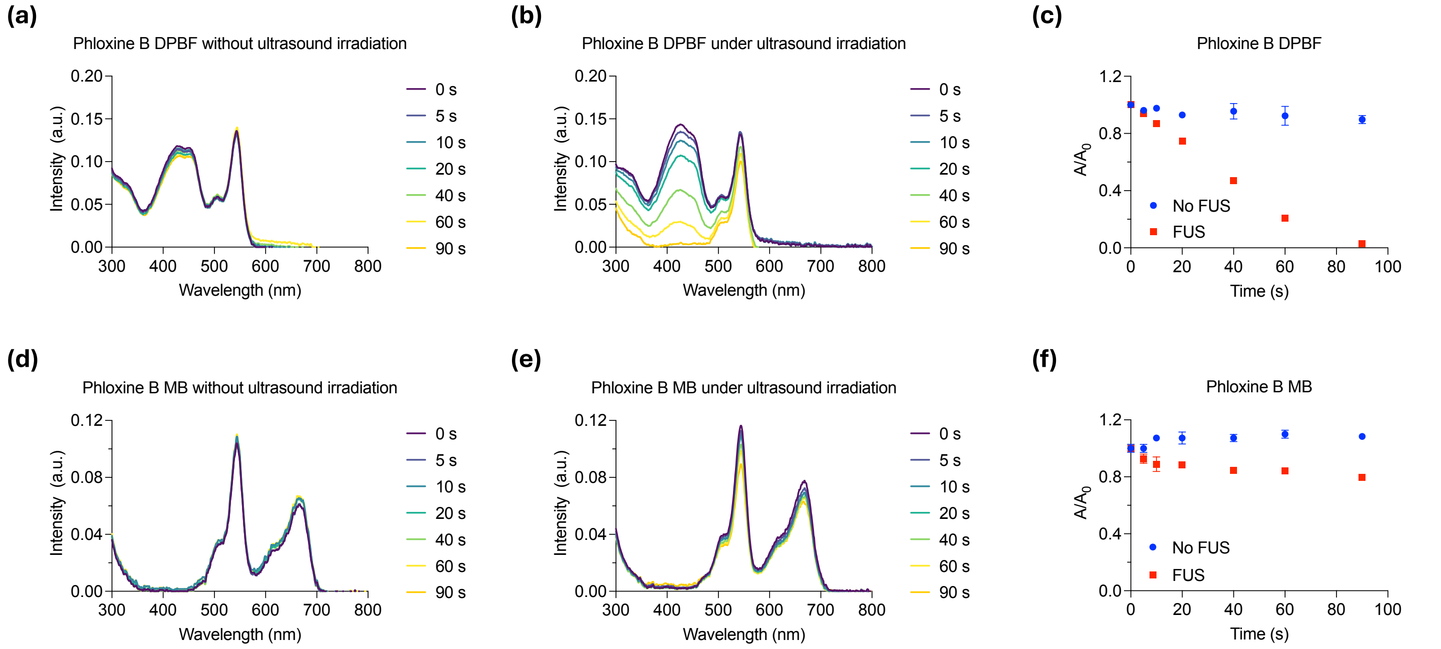


**Figure S14**. The ultrasound-induced ROS generation of sonosensitizer Phloxine B. (a) UV-Vis spectra showing no significant change in DPBF absorption in the presence of Phloxine B without ultrasound stimulation (1.5 MHz, 1.5 MPa, pulse 500 ms on, 500 ms off). (b) Time-dependent UV-Vis spectra demonstrating DPBF decomposition and $^{1}O_2$ generation by Phloxine B under ultrasound stimulation. (c) Quantitative analysis of DPBF decomposition induced by ultrasound in the presence of Phloxine B compared to controls (n > 3 per group). (d) UV-Vis spectra indicating negligible MB decomposition in the absence of ultrasound. (e) Time-dependent UV-Vis spectra demonstrating MB degradation by •OH produced from Phloxine B ultrasound stimulation. (f) Quantitative analysis of MB decomposition with and without ultrasound irradiation in the presence of Phloxine B (n > 3 per group). All plots show mean ± SEM unless otherwise mentioned.

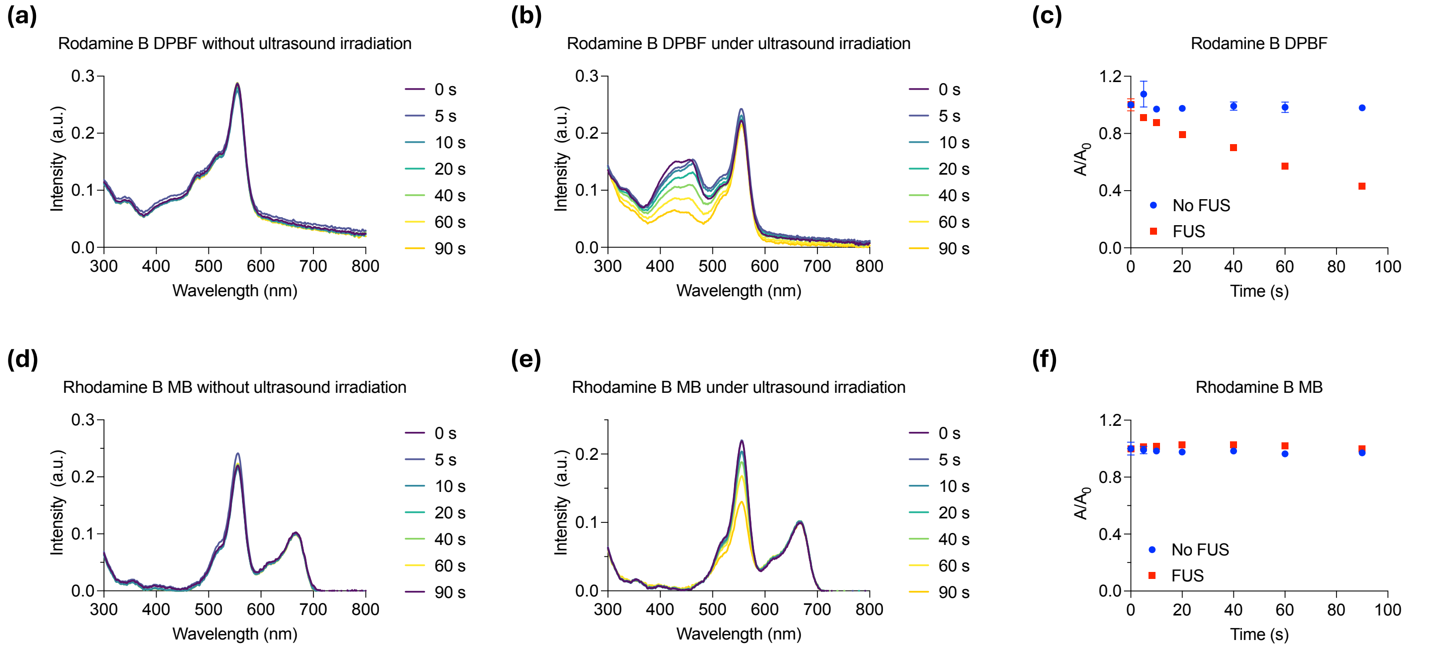


**Figure S15**. The ultrasound-induced ROS generation of sonosensitizer Rhodamine B. (a) UV-Vis spectra showing no significant change in DPBF absorption in the presence of Rhodamine B without ultrasound stimulation (1.5 MHz, 1.5 MPa, pulse 500 ms on, 500 ms off). (b) Time-dependent UV-Vis spectra demonstrating DPBF decomposition and $^1O_2$ generation by Rhodamine B under ultrasound stimulation. (c) Quantitative analysis of DPBF decomposition induced by ultrasound in the presence of Rhodamine B compared to controls ($n > 3$ per group). (d) UV-Vis spectra indicating negligible MB decomposition in the absence of ultrasound. (e) Time-dependent UV-Vis spectra demonstrating MB degradation by •OH produced from Rhodamine B ultrasound stimulation. (f) Quantitative analysis of MB decomposition with and without ultrasound irradiation in the presence of Rhodamine B ($n > 3$ per group). All plots show mean ± SEM unless otherwise mentioned.

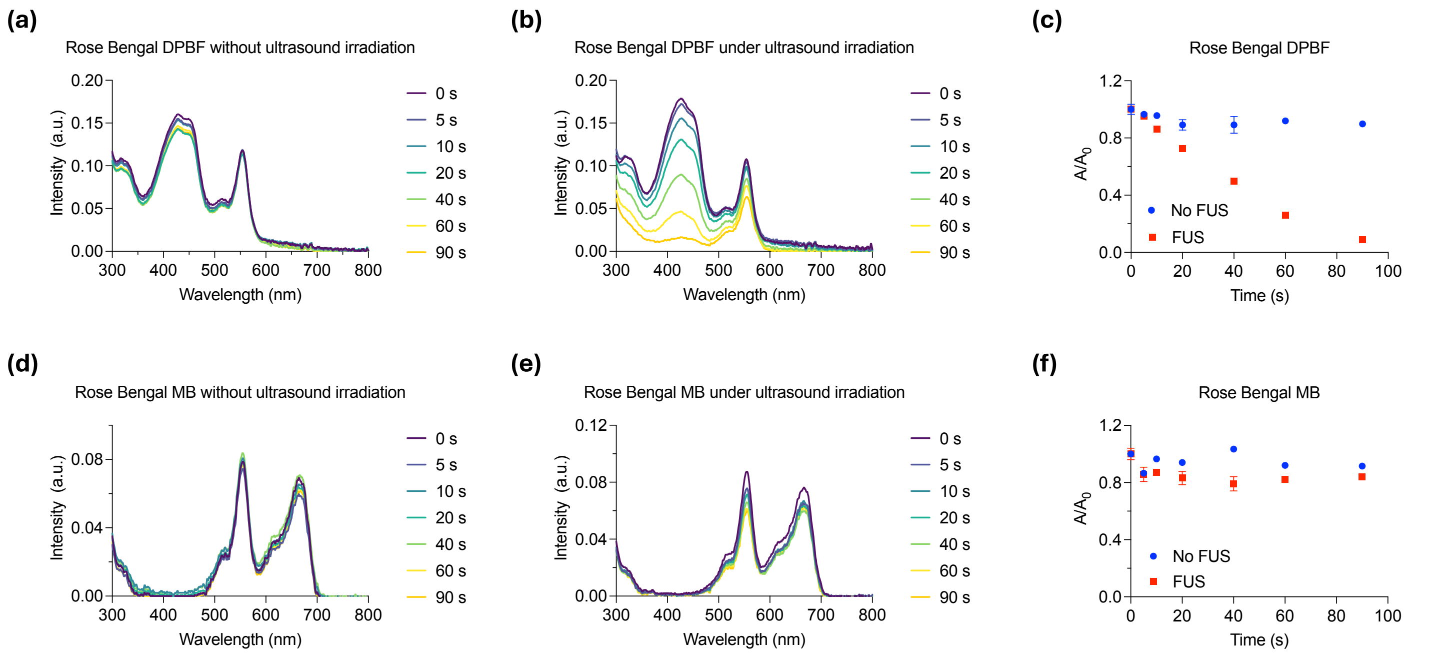


**Figure S16**. The ultrasound-induced ROS generation of sonosensitizer Rose Bengal. (a) UV-Vis spectra showing no significant change in DPBF absorption in the presence of Rose Bengal without ultrasound stimulation (1.5 MHz, 1.5 MPa, pulse 500 ms on, 500 ms off). (b) Time-dependent UV-Vis spectra demonstrating DPBF decomposition and $^1O_2$ generation by Rose Bengal under ultrasound stimulation. (c) Quantitative analysis of DPBF decomposition induced by ultrasound in the presence of Rose Bengal compared to controls ($n > 3$ per group). (d) UV-Vis spectra indicating negligible MB decomposition in the absence of ultrasound. (e) Time-dependent UV-Vis spectra demonstrating MB degradation by •OH produced from Rose Bengal ultrasound stimulation. (f) Quantitative analysis of MB decomposition with and without ultrasound irradiation in the presence of Rose Bengal ($n > 3$ per group). All plots show mean ± SEM unless otherwise mentioned.
